# Population genetics reveals divergent lineages and ongoing hybridization in a declining migratory fish species complex

**DOI:** 10.1101/2021.12.04.471201

**Authors:** Quentin Rougemont, Charles Perrier, Anne-Laure Besnard, Isabelle Lebel, Yann Abdallah, Eric Feunteun, Elodie Réveillac, Emilien Lasne, Anthony Acou, David José Nachón, Fernando Cobo, Guillaume Evanno, Jean-Luc Baglinière, Sophie Launey

**Author notes:** present address (Quentin Rougemont, Elodie Réveillac, Emilien Lasne).

## Abstract

Deciphering the effects of historical and recent demographic processes responsible for the spatial patterns of genetic diversity and structure is a key objective in evolutionary and conservation biology. Using population genetic analyses, we investigated the demographic history, the contemporary genetic diversity and structure, and the occurrence of hybridization and introgression of two species of anadromous fish with contrasting life history strategies and which have undergone recent demographic declines, the allis shad (*Alosa alosa*) and the twaite shad (*Alosa fallax*). We genotyped 706 individuals from 20 rivers and 5 sites at sea in Southern Europe at thirteen microsatellite markers. Genetic structure between populations was lower for the nearly semelparous species *A. alosa*, which disperses greater distances compared to the iteroparous species, *A. fallax*. Individuals caught at sea were assigned at the river level for *A. fallax* and at the region level for *A. alosa*. Using an approximate Bayesian computation framework, we inferred that the most likely long term historical divergence scenario between both species and lineages involved historical separation followed by secondary contact accompanied by strong population size decline. Accordingly, we found evidence for contemporary hybridization and bidirectional introgression due to gene flow between both species and lineages. Moreover, our results support the existence of at least one distinct species in the Mediterrannean sea: *A. agone* in Golfe du Lion area, and another divergent lineage in Corsica. Overall, our results shed light on the interplay between historical and recent demographic processes and life history strategies in shaping population genetic diversity and structure of closely related species. The recent demographic decline of these species’ populations and their hybridization should be carefully considered while implementing conservation programs.

## Introduction

Reconstructing how historical and recent processes shape present genetic diversity is an important step in evolutionary biology. In particular, long-term processes such as vicariance events, long term contraction during the last ice age or postglacial recolonization are well known to shape current patterns of genetic diversity (Hewitt 1996). In addition, several species and/or populations are currently declining at a fast pace due to human activity or climate change (Ceballos et al. 2020, Ryan et al. 2018). The interplay of these processes may leave counter-intuitive signatures on contemporary patterns of genetic diversity, making it difficult to decorrelate them and quantify their respective roles. In addition, populations within species or closely related species may differ in terms of life history traits, which can result in contrasting levels of population genetic structure and diversity.

In particular, the movement of species or dispersal processes are key factors affecting genetic structure through space and time (Cayuela et al. 2018). Many migratory species have undergone sharp declines in population sizes worldwide (Wilcove and Wikelski 2008; Limburg and Waldman 2009), which can have evolutionary consequences for these species. Indeed, small populations can display higher rates of genetic drift, accumulation of deleterious alleles and loss of genetic variability, which can ultimately threaten their genetic integrity and adaptive potential (Frankham 2005). Therefore, quantifying a species’ evolutionary potential is vital for the conservation of wild and managed populations. This task can benefit from reconstruction of demographic history from genetic data (Rougemont et al. 2020), monitoring of population genetic parameters including genetic diversity, effective population size (*Ne*), and quantifying hybridization between closely related groups or species (Barton and Hewitt 1985) as well as their positive (Abbott et al. 2013) or negative consequences on hybrid fitness (Mikkelsen and Irwin 2021).

Diadromous fish are keystone species that move between freshwater and marine environments. Many of these species, including salmonids, eels, sturgeons or shads, have undergone steep declines in their population size due to human activities such as dams building, over-harvesting, habitat degradation, and pollution (Parrish et al. 1998; Waters et al. 2000; Limburg and Waldman, 2009). Some of them (e.g., salmonids and shads) tend to return to their natal river for breeding (i.e “homing” behavior), as demonstrated by otolith analysis (Tomás et al. 2005; Walther and Thorrold 2008; Perrier et al. 2011; Martin et al. 2015; Randon et al. 2017). This homing behavior fosters local adaptation, which is beneficial in stable environmental conditions (Keefer and Caudill 2014), but can also isolate some populations and result in smaller local effective population sizes. Consequently, in the context of environmental changes, this strategy may lead to less stable populations as compared to those displaying higher levels of gene flow and effective population size (Frankham 1997, 2005), although this question has given rise to recent debates (e.g. Kyriazis et al. 2020; Teixeira & Huber et al. 2021; Garcia-Dorado et al. 2021).

The allis shad, *Alosa alosa* (Linnaeus, 1758), and the twaite shad, *Alosa fallax* (Lacépède, 1803), are two closely related anadromous *Clupeidae* species that have undergone steep declines in abundance and distribution range (Baglinière et al. 2003a). Since the middle of the 20^th^ century, declines for both species’ have mostly been attributed to freshwater habitat degradation inducing loss of spawning grounds and over-harvesting (Aprahamian et al. 2003; Limburg and Waldman 2009). Both species are now mostly restricted to large rivers in Portugal, Spain, France and the United Kingdom (mainly *A. fallax*). shows Presently, this decline appears more substantial for *A. alosa;* according to the last assessment of IUCN status for France, *A. alosa* is now considered as critically endangered (Baglinière et al. 2020a), while *A. fallax* is considered as vulnerable (Baglinière et al. 2020b). It is expected that this decrease in abundance has had important repercussions on the genetic diversity of both species. Importantly, a decrease in census population size will reduce effective population size and decrease selection efficacy, but the effects on either species may differ given differences in life history strategies despite their close phylogenetic relationship (Faria et al. 2012). For example, *A. alosa* is nearly-semelparous while *A. fallax* is iteroparous (Mennesson-Boisneau et al. 2000) and *A. alosa* displays a stronger dispersal behavior than *A. fallax* (Taverny and Elie 2001; Jolly et al. 2012; Martin et al. 2015; Nachón et al. 2020). Following the study of Hasselman et al. (2013) we predict that iteroparity sets a constraint on *A. fallax* dispersal and migratory distance due to stronger selection for homing every year for breeding (Jolly et al. 2012). In contrast, longer migratory distance may facilitate gene flow in the semelparous *A. alosa* (Martin et al. 2015; Randon et al. 2017).

Because of these differences in dispersal and parity mode, we expected stronger gene flow and weaker population genetic structure in *A. alosa* compared to *A. fallax*. The genetic distinctiveness of *A. alosa* and *A. fallax* has been confirmed (Faria et al. 2004; Alexandrino et al. 2006; Jolly et al. 2012), but the details of their demographic history of prior isolation and subsequent contact has not been fully resolved (Faria et al. 2012; Taillebois et al. 2020). This issue could benefit from demographic modeling based on genetic data. Such an approach would help to decipher whether these species evolved separately before a secondary contact, or whether they could have evolved in sympatry, which has important implications for understanding speciation processes, as well as for the conservation of the species. This might especially influence the rate at which hybridization could result in a generalized meltdown of both species into one. In fact, hybrids between both species have been documented in some geographic areas like Portugal, United Kingdom and France (Alexandrino et al. 2006; Coscia et al. 2010; Jolly et al. 2011; Faria et al. 2012; Taillebois et al. 2020), suggesting that barriers to gene flow between species are permeable. In the context of river fragmentation, barriers (e.g. dams) generate “forced” common spawning grounds between *A. alosa* and *A. fallax* (Alexandrino et al. 2006), which would otherwise be spatially segregated along the river network (Mennesson-Boisneau et al. 2000). It is therefore expected that both species hybridize frequently in rivers with high levels of fragmentation (see also Taillebois et al. 2020). In addition, the taxonomic status of *A. fallax* within the Mediterranean Sea is debated (Chiesa et al. 2014; Baglinière et al. 2020c), despite genetic and morphological support for the existence of two lineages (Le Corre et al. 2005; Bianco, 2005). Analysis of a broader geographic sample such as that described herein may help to resolve this taxonomic conundrum.

In this study we used 13 microsatellite markers (Rougemont et al. 2015) to *i*) determine the extent of genetic diversity and structure among *Alosa alosa* and *Alosa fallax* along the Atlantic and Mediterranean coasts, *ii*) determine the historical process of divergence between species using approximate Bayesian computations, *iii*) contrast the patterns of long-term historical gene flow between species and among major genetic groups versus contemporary dispersal at sea in each species, and *iv*) test the power of this marker set to assign individuals captured at sea. In doing so, we tested the hypotheses that *i*) both species were formerly isolated and came into secondary contact, *ii*) both species differ in their genetic structure, with the more dispersive, semelparous species showing weaker genetic structure than the iteroparous species, *iii*) hybrids are frequently found, potentially as a result of frequent “forced” common spawning grounds and *iv*) individuals caught at sea can be confidently assigned to rivers or geographic region, depending on the species.

## Materials and Methods

### Study area and sampling

A total of 706 individuals from both species (367 *A. alosa* and 339 *A. fallax*) were sampled from 20 rivers distributed along the French Atlantic coast (N=514), Spanish coast (N= 56 *A. fallax*), Mediterranean coast (N=92 *A. fallax*) and Corsica (N= 65 *A. fallax*) A subset of these (40 *A. alosa* and 65 *A. fallax*) were captured at sea along the Atlantic coast and were also used for population assignment (**Fig. 1, Table1** and **Table S1**). Individuals collected in rivers were captured between 2009 and 2012 either by nets or trapping and a fin clip was stored in 95% ethanol, while individuals captured at sea were collected by professional fishermen and identified phenotypically. Scales were also collected from carcasses after breeding migration and conserved in paper envelopes. Collections of scales (stored at INRAE Rennes, France) from cohorts 1997 to 1999 and 2001 (Aulne and Charente rivers) were also used (**Table1**).

**Figure 1:**
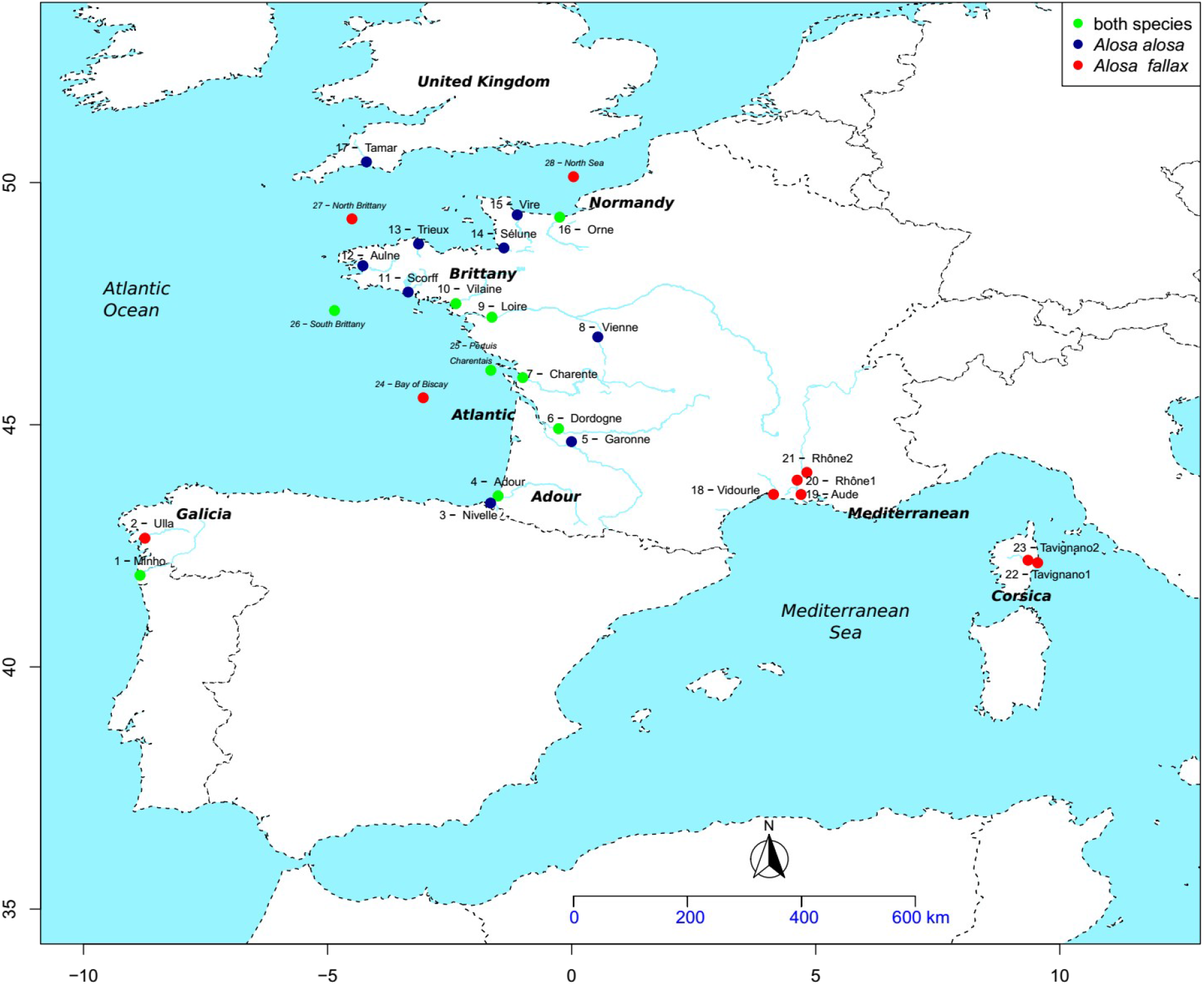
Map showing the location of sampling sites across France, United-Kingdom and Spain. Sample site numbers correspond to those given in table 1.

**Table 1:**
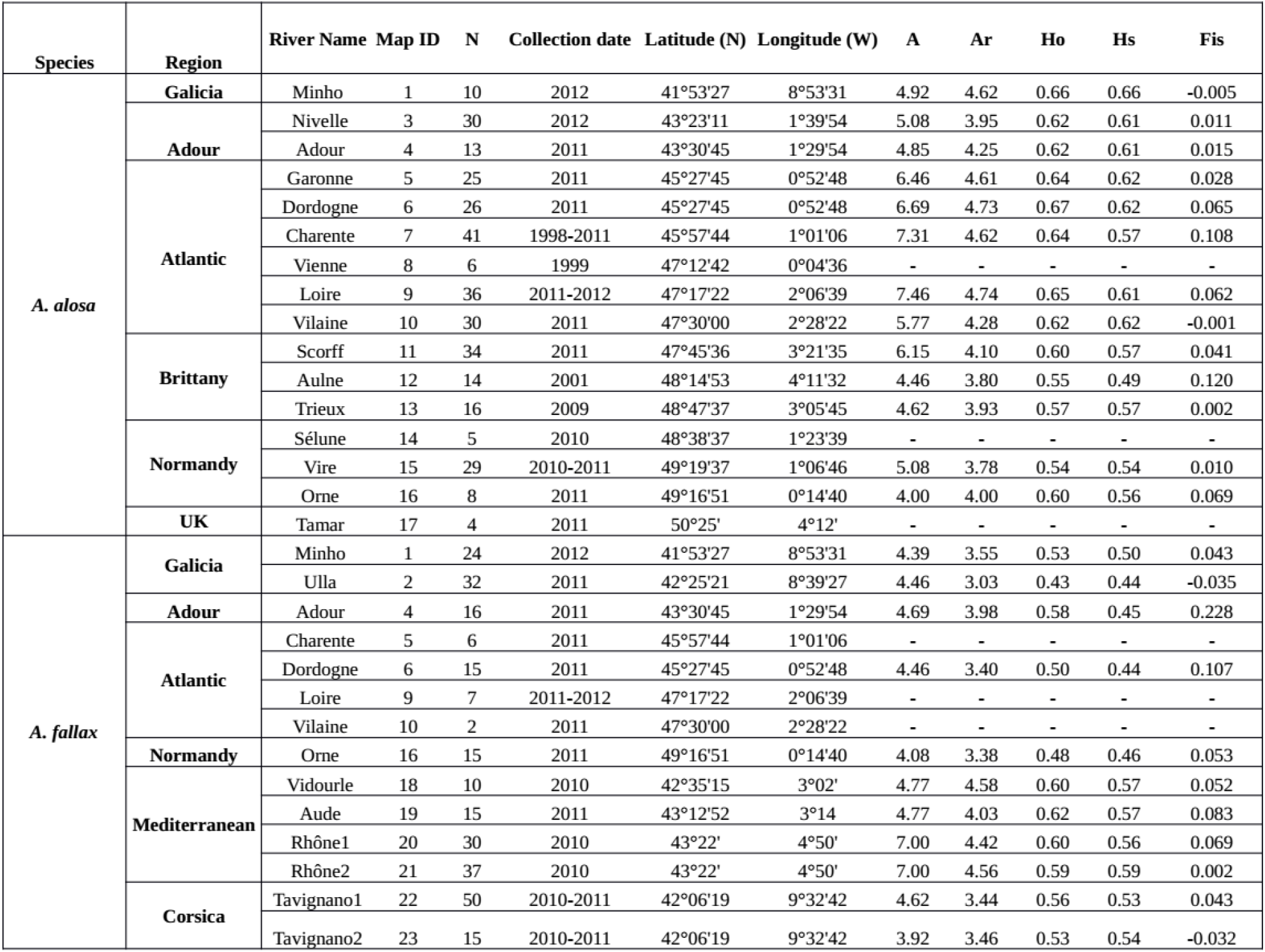
Sampling information for each species. N = sample Size. Genetic diversity indices are as follows; A: number of allele. Ar: Allelic richness adjusted for sample size (8 in A. alosa. 15 in A. fallax). Ho: Observed Heterozygosity. He: Unbiased expected heterozygosity. Fis: Coefficient of inbreeding. Details of shad populations captured at sea are available in the supplementary material (Table S1)

### Molecular methods

Genomic DNA was extracted using a Chelex protocol (modified from Estoup et al. 1996). We genotyped each individual at 13 microsatellite loci specifically developed for *A. alosa* and *A. fallax*(Rougemont et al. 2015). Occurrence of null alleles and scoring errors due to large allelic dropout or stuttering were checked using Micro-Checker v2.2.3 (Oosterhout et al. 2004). Linkage disequilibrium (LD) was inspected using Genepop 4.0 (Rousset 2008).

#### Genetic diversity and HWE within populations

Fish captured at sea for which the river of origin was unknown and hybrids were excluded from genetic diversity and Hardy-Weinberg equilibrium (HWE) analyses, leaving a total of 586 fish (315 *A. alosa* and 271 *A. fallax*). For each population (sample in each river) of each species, diversity indices (observed heterozygosity (Ho) and expected Heterozygosity (He) under HWE) were calculated with Genetix 4.0.5 (Belkhir et al. 1996). Significance of heterozygosity comparisons was assessed with a Wilcoxon sign rank test. Deviations from HWE and linkage disequilibrium were tested for each locus within each population and globally wiht Genepop 4.0 (Rousset 2008). Fstat version 2.9.3 (Goudet 1995) was used to calculate inbreeding coefficient (Fis), allele number, and allelic richness (Ar), with Ar differences between species tested using 4,000 permutations. We tested the significance of the differences in genetic diversity among each river within species using a linear mixed model. First we tested the effect of the “River” factor considered as a fixed effect on levels of Allelic Richness considered as the response variable. We included the locus as a random effect to account for their variation and for their random representation of the genomic variability. An Anova was used to test the effect of the River and the amount of variance was quantified using the conditional and marginal coefficient of determination (R2c and R2m respectively describing the variance accounted for by locus variability and by “River” alone. Finally differences among rivers were tested using TukeyHSD post-hoc comparisons on the models. Tests were implemented using the lme4 package (Bates et al. 2015), MuMIn (Barton, 2021) and multComp (Hothorn et al. 2008)

### Hybrid identification

We were interested in accurately distinguishing purebred individuals from hybrids. We used a similar framework to the one of Vähä and Primmer (2006) using Structure 2.3.3 (Pritchard et al. 2000) and NewHybrids 1.1 (Anderson and Thompson 2002). For the purpose of hybrid identification we assumed K=2 where each cluster is composed of a single species contributing to the total gene pool of the sample. Structure was used without prior information about the species classification using an admixture model with (*i*) correlated allele frequency (Falush et al. 2003) and (*ii*) 500,000 burn-in steps followed by 1,000,000 MCMC iterations, replicated 10 times. Individual admixture proportions (q-values) and their 90% confidence intervals were averaged over the ten replicates and used to assign individuals to their respective genetic clusters. NewHybrids assumes that the sample is drawn from a mixture of pure individuals and hybrids (F1, F2 or backcrosses) so that the q-value inferred with this method is a discrete variable (Anderson and Thompson, 2002). NewHybrids was used to calculate the posterior probability for an individual to belong to one of the following classes: purebred *A. alosa*, purebred *A. fallax*, hybrid F1, hybrid F2 or backcross. The q-values were summed over all hybrid categories (F1, F2 and backcross) because preliminary tests showed that performance was greater under these conditions. Hence, hybrids, regardless of their hybrid class, were distinguished from purebreds. Uniform priors were used for allele frequency and admixture estimations were performed with a burn in of 1,000,000 steps followed by 1,000,000 iterations. Tests using Jeffrey priors, instead of uniform, yielded similar results. Results are based on the average of 10 runs performed with random seed.

### Performance of Admixture Analysis

To test the correctness of assignment of Structure and NewHybrids, the software HybridLab (Nielsen et al. 2006) was used. Individuals showing q-value >0.9 with both methods were chosen randomly and 3,500 simulated genotypes of each parental species were generated and divided into 10 datasets of 350 individuals. The sample size was chosen to be close to our real dataset. Then 10 other datasets were created containing both parental and hybrid individuals (F1, F2, backcrosses). Ten individuals of each hybrid category were incorporated into the datasets of 350 individuals to represent the actual sample size of our empirical dataset and to incorporate a small proportion of hybrids. The simulated dataset was analyzed with Structure and NewHybrids to calculate (1) the hybrid proportion, which is the number of individuals classified as hybrid divided by the total number of individuals in the sample, (2) efficiency, (3) accuracy and (4) overall performance of these methods following the definition of Vähä and Primmer (2006). We used a q-value threshold of 0.90 for hybrid classification.

We tested the significance of the differences in genetic diversity among each river within species using a linear mixed model. First we tested the effect of the “River” factor considered as a fixed effect on levels of Allelic Richness considered as the response variable. We included the locus as a random effect to account for their variation and for their random representation of the genomic variability. An anova was used to test the effect of the River and the amount of variance was quantified using the conditional and marginal coefficient of determination (R2c and R2m respectively describing the variance accounted for by locus variability and by “River” alone, Nakagawa and Schielzeth 2013). Finally differences among rivers were tested using TukeyHSD post-hoc comparisons on the models. Tests were implemented using the lme4 package (Bates et al. 2015), MuMIn (Barton, 2021) and multComp (Hothorn et al. 2008)

### Population genetic structure

Genetic structure was assessed excluding hybrids (based on their genotype) and fish captured at sea. The extent of genetic structure was quantified using the pairwise *F_ST_* estimator θ_ST_ of (Weir and Cockerham 1984) between sample pairs with Fstat, and with significance assessed using 10,000 permutations. We tested for a signal of isolation by distance (IBD) in our data in both species separately using a mantel test on matrices of linearized *F_ST_* using *F_ST_*/(1 – *F_ST_*) against the logarithm of the waterway distances measured between river mouths manually in ArcGIS following the coastline. We next plotted the signal of isolation by distance (Fig 3a) in each species separately with the ggplot2 package and tested the strength of the relationship (R^2^) and its significance using a simple linear model.

The chord distance (Cavalli-Sforza and Edwards 1967) was used to quantify genetic differentiation between sample pairs and to construct neighbor-joining phylograms (Saitou and Nei 1987) with MSA 4.05 (Dieringer and Schlötterer 2003). Trees were computed using the software Phylip 3.6 using a maximum likelihood optimality criterion (Felsenstein 1995) and 10,000 permutations were conducted to establish bootstrap support for the nodes. The results were then visualized using Tree view 1.6 (Page, 1996). Individual genetic clustering was further investigated without *a priori* definition of population boundaries using STRUCTURE. The admixture and correlated allele frequency model was used to detect a number of genetic clusters (K) varying from 1 to 14 separately for *A. alosa* and *A. fallax*. Fifteen replicates were run for each K with a burn-in period of 400,000 followed by 400,000 MCMC iterations. The optimal number of clusters was evaluated using the likelihood distribution (Pritchard et al. 2000) and the ΔK method (Evanno et al., 2005). Results were combined with Structure harvester (Earl and vonHoldt 2012) and graphs were plotted in R using Pophelper (Françis 2017). Finally, we used a multivariate method, the Discriminant Analysis of Principal Components implemented in the Adegenet Package (Jombart 2008) in R (R Development Core Team 2012), to inspect structure between populations for each species. We used the function find.cluster, which uses k-means to find the optimal number of clusters and selected the number of groups with the lowest bayesian information criteria (BIC). We also used the alpha score to choose the optimal number of principal components to retain.

### Genetic stock assignment

Genetic assignment of fish captured at sea was performed using three different methods. First Structure was run with fish captured at sea included in the dataset. The same settings as described above were used. Second, DAPC assignments were performed with fish from the sea as supplementary individuals using the previously defined parameters. Third, we used the Bayesian method implemented in GeneClass 2 (Piry et al. 2004) specifically designed for assignment tests. The likelihood that any fish captured at sea came from one of the sampled populations was tested using the resampling algorithm described by Paetkau et al. (2004) with 100,000 simulated individuals. Fish that displayed a probability <0.01 were assumed to come from an un-sampled reference population and were excluded from the assignment tests.

### Demographic history: approximate Bayesian computations

The demographic history of divergence between *A. alosa* and *A. fallax*, as well as between *A. fallax* from Atlantic vs *A. fallax* from the Mediterranean sea, was investigated using an ABC framework. To avoid any bias due to sparse sampling, intra population structure or isolation by distance (e.g. Mason et al. 2020), we focused our between species comparison on *A. fallax* and *A. alosa* sampled along the Atlantic coast. Our between lineage comparison included all individuals from the Rhone river and all individuals from Corsica.

We excluded all putative hybrids to avoid favoring models of ongoing gene flow because our focus was on long-term patterns of gene flow. A total of four scenarios of divergence were compared (**Fig S1**). The model of strict isolation (SI) assumes that an ancestral population of size *N_ANC_* splits instantaneously at time *T_split_* into two daughter populations of constant and independent size *N*_pop1_ and *N*_pop2_. The split is not followed by any gene flow and the population can either undergo an instantaneous bottleneck or an expansion at *T_split_*. In contrast, the three other models assume various rates of gene flow following the instantaneous split. This migration occurs at a rate *M* = 4 *N_0_.m.* with M_1←2_ being the number of migrants from population 2 to population 1 and M_2←1_ being the reciprocal. In the model of ancestral migration (AM) the first generations of divergence are followed by gene flow until time *T_am_* (**Fig S1**), at which point there will be no further gene flow (going forward in time). In the model of isolation with migration (IM) genes flow occurs continuously from *T_split_* to the present and at a constant rate each generation. Under the secondary contact (SC) model, *T_split_* is followed by a period of strict isolation and then by a period of secondary contact at *T*_sc_ generations ago that is still ongoing. We used uniform priors for model choice and parameter estimation. Simulation of microsatellites strictly followed the procedure of Illera et al. (2014) and later modified by Rougemont et al. (2016). In detail, the ms software (Hudson, 2002) was used to perform coalescent simulations under an infinite-site model of mutation. Binary simulated data from ms were converted into microsatellite data using a generalized stepwise mutation model (GSM) in which the probability of changes of the repeat number in each mutation event was modeled by a geometrical parameter α following a uniform prior distribution sampled on the interval 0–0.5. Each effective population size *N_pop1_, N_pop2_, N_ANC_* was scale by the parameter θ = 4*N_ref_* *μ, with *N_ref_* representing the effective population size of an arbitrarily chosen reference population (*N_ref_*, here set to 50,000) and μ representing the mutation rate per generation (chosen as μ = 2.5e^-4^ bp/generations). We chose μ according to values frequently observed in fishes (Shimoda et al.,1999; Yue, David & Orban, 2006). All divergence time parameters (T_split_, T_sc_, T_am_) were also scaled by the parameter 4*N_ref_*. Priors for the effective population size were set to [0-500,000], [0-2,500,000] for the ancestral population size and [0-1,000,000] generations for the split time **(table S7)**. These correspond to large and uninformative priors following Cornuet et al. (2010) representing a set of biologically realistic values given the history of glacial cycles in Europeans biomes (e.g. Hewitt, 1996).

Given *i*) the arbitrarily fixed mutation rate and *ii*) the interspecific variability in age at maturity, we did not attempt to convert the inferred divergence time or timing of secondary contact into a number of years to avoid over-interpretation of these parameters. Yet, the ratios of divergence time or symmetry in effective population size between species remained biologically relevant since the effect of the assumed mutation rates cancels out. All computations took into account differences in sample size for each of the thirteen loci. four millions simulations composed of the thirteen microsatellite loci were computed under each demographic model.

Summary statistics were computed from the transformed microsatellite data and included the average and standard deviation values of: the number of alleles (A), Allelic richness (Ar), observed and expected heterozygosity (Ho and He, respectively), allele size in base pairs, the Garza-Williamson index (GW, Garza & Williamson, 2001), G_ST_ (Nei 1973) and δμ2 (Goldstein et al., 1995). All statistics were computed using R scripts (R Development Core Team, 2015).

#### Model selection

We evaluated the posterior probabilities of each demographic model using an ABC framework implemented in the abc package in R (Csilléry et al., 2012). We computed posterior probabilities using a feed forward neural network based on a nonlinear conditional heteroscedastic regression in which the model is considered as an additional parameter to be inferred. In the rejection step, we retained the 0.02% of simulations closest to the observed summary statistics, which were subsequently weighted by an Epanechnikov kernel. The regression step was performed using 50 neural networks and 15 hidden layers.

#### Parameter estimation and cross-validation

Parameter estimation was performed for the best models using nonlinear regressions. We first used a logit transformation of the parameters on the 4,000 best replicate simulations providing the smallest Euclidean distance to the observed data (Csilléry, et al. 2012). We then jointly estimated the posterior probability of each parameter using the neural network procedure implemented in the abc package. We obtained the best model by weighted nonlinear regressions of the parameters on the summary statistics using 50 feed-forward neural networks and 15 hidden layers. Cross-validation was performed by computing the robustness of the model choice using a total of 4,000 pseudo observed datasets (PODS) sampled randomly from each model. The same ABC procedure as for the empirical dataset was performed but this time considering a given model M instead of the empirical data. Then the robustness between two given models *M_1_* and *M_2_* was computed as:

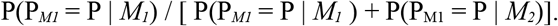

Where P(P_*M1*_ = P | *M_1_*) represents the probability of correctly supporting *M1* given the observed posterior probability *P*, and P(P_M1_ = P | *M2*) is the probability of erroneously supporting *M_1_* given that the true model is M2 (Fagundes et al., 2007). Parameter estimation was performed for the best model by running another round of simulations for a total of 4 million simulations. The whole pipeline used for ABC can be found at: https://github.com/QuentinRougemont/MicrosatDemogInference.

## Results

A total of 706 individuals were genotyped at 13 microsatellite loci and 180 different alleles were found, with 153 in *A. alosa* and 138 in *A. fallax*. No evidence of null alleles was detected. In *A. alosa* one test of linkage disequilibrium (LD) out of 936 comparisons was significant. Similarly, one LD test out of 858 comparisons was significant in *A. fallax*.

### Genetic diversity

Fis was significant for only one marker (Alo29) in the Loire River population of *A. alosa*. All other markers across all populations did not show deviations from HWE or significant Fis values. The total number of alleles varied from 4 to 20 in *A. alosa* and from 4 to 17 in *A. fallax*. The two species shared 62% of alleles. Mean adjusted allelic richness (*Ar*) per population ranged from 3.74 to 5.94 in *A. alosa* and from 3.44 to 5.70 in *A. fallax* (**Table 1** for details by river and **Table S2, Table S3** for details for each river and each marker). Mean observed heterozygosity was 0.596 (range: 0.49–0.62) in *A. alosa* and 0.523 (range: 0.44–0.59) in *A. fallax* and differed significantly between species (P<0.005, 5,000 permutations). In contrast, the difference in *Ar* between species was not significant (P=0.061, 5,000 permutations). For *A. alosa, Ar* and *He* were greatest in the Dordogne, Minho and Loire River and lowest in the Vire, Aulne and Trieux rivers. Accordingly, our linear models indicated a significant effect of the River factor (p = 0.0010**, **Table S4**, R2m = 0.065, R2c = 0.700). Yet, none of the observed differences between rivers were significant according to our TukeyHSD comparisons (p > 0.05, **Table S5**). For *A. fallax, Ar* and *He* were highest in the Rhône (Mediterranean) and lowest in the Ulla, Orne and Dordogne (Atlantic). Accordingly, our linear models indicated a significant effect of the River factor (p = 8e-5***, Table S4, R2m = 0.147, R2c = 0.496, Table S6). In this case the Ulla was the only population with significantly less allelic richness (p < 0.01) than the Rhône. None of the remaining comparisons were significant (Table S5).

#### Hybrid identification

Results of admixture analyses in Newhybrids and Structure using simulated genotypes from Hybridlab varied depending on the presence or absence of hybrids (Table S6). Both Structure and Newhybrids produced some false hybrids in the purebreds’ dataset and this overestimation was greater in Structure than in Newhybrids (Table S6). In brief, both software displayed an accuracy and efficiency at a q-value threshold of 0.90, although NEWHYBRIDS displayed a slightly higher efficiency and accuracy compared to STRUCTURE with q > 0.90. In brief, NewHybrids displayed slightly higher efficiency and accuracy at a q-value threshold of 0.90 in the presence of hybrids with both software displaying accuracy and efficiency above 0.90.

Regarding our empirical data, both species were separated in two fully distinct clusters with average q-values greater than 0.99 for *A. alosa* and *A. fallax* with both Structure and NewHybrids. Both methods reclassified 13 individuals captured at sea that were morphologically identified as A. alosa but genotypically classified as *A. fallax* and 1 individual wrongly identified as *A. fallax* but genotypically classified as *A. alosa*. A total of 25 individuals displayed a q-value less than 0.9 with either of the methods and were not classified as purebreds. All hybrids identified by NewHybrids had a q-value greater than 0.8 and were all identified as hybrids in Structure (Table S7). From a geographic standpoint, the majority (80%) of the 25 individuals came from three major areas: namely the Charente River and Pertuis Charentais (28%), the Loire River (28%) and South Brittany/Scorff River (24%).

### Population genetic structure

Global inter-species *F*_ST_ was 0.240 (95% CI: 0.194 – 0.286) and significant (P<0.00087. 15,000 permutations). Global *F*_ST_ for *A. alosa* was 0.046 (95% CI: 0.034 – 0.058) (P<0.0001. 10,000 permutations). Global *F*_ST_ for *A. fallax* was higher with a value of 0.219 (95% CI: 0.175 – 0.263) (P<0.0001. 10,000 permutations). In *A. alosa*, 89% of pairwise *F*_ST_ comparisons were significant after Bonferroni corrections (**Table 2**) compared to 82% in *A. fallax*. Levels of differentiation between sampling localities were lower in *A. alosa* than in *A. fallax*. Populations sampled from the same river basin (i.e Rhône, or Tavignano) were not significantly differentiated from each other (**Table 2),** but were significantly differentiated from all other rivers. Similarly, Mediterranean *A. fallax* populations (Aude, Rhône, Vidourle) were not differentiated among one another but were strongly differentiated from Atlantic populations. The Tavignano (Corsica) population was significantly differentiated from all other rivers (*F_ST_* ranged between 0.233 and 0.358). *A. fallax* populations from the Minho, Ulla and Orne were also distinct from all other populations (*F_ST_* ranged between 0.08 and 0.289), but comparisons with the Tavignano were the highest (**Table 2, Table S8** for interspecific comparison).

**Table 2:** pairwise F_ST_ estimates for each species (non significant values are in bold italic, samples are sorted from South to North)

**A. fallax**
|  | Minho | Ulla | Adour | Dordogne | Orne | Aude | Rhône1 | Rhône2 | Vidourle | Tavignano |
| --- | --- | --- | --- | --- | --- | --- | --- | --- | --- | --- |
| Ulla | 0.147 |  |  |  |  |  |  |  |  |  |
| Adour | 0.081 | 0.204 |  |  |  |  |  |  |  |  |
| Dordogne | 0.088 | 0.233 | 0.029 |  |  |  |  |  |  |  |
| Orne | 0.207 | 0.289 | 0.180 | 0.144 |  |  |  |  |  |  |
| Aude | 0.224 | 0.253 | 0.169 | 0.217 | 0.170 |  |  |  |  |  |
| Rhône 1 | 0.229 | 0.253 | 0.182 | 0.220 | 0.173 | **0.010** |  |  |  |  |
| Rhône 2 | 0.227 | 0.264 | 0.184 | 0.225 | 0.166 | **0.011** | **0** |  |  |  |
| Vidourle | 0.245 | 0.289 | 0.177 | 0.240 | 0.217 | **0.011** | **0.001** | **0.016** |  |  |
| Tavignano1 | 0.286 | 0.328 | 0.253 | 0.288 | 0.323 | 0.233 | 0.237 | 0.252 | 0.242 |  |
| Tavignano2 | 0.303 | 0.345 | 0.260 | 0.308 | 0.358 | 0.248 | 0.249 | 0.268 | 0.262 | **0.011** |

**A. alosa**
|  | Minho | Nivelle | Adour | Charente | Dordogne | Garonne | Loire | Vilaine | Scorff | Aulne | Trieux | Vire |
| --- | --- | --- | --- | --- | --- | --- | --- | --- | --- | --- | --- | --- |
| Nivelle | 0.119 |  |  |  |  |  |  |  |  |  |  |  |
| Adour | 0.094 | 0.060 |  |  |  |  |  |  |  |  |  |  |
| Charente | 0.058 | 0.093 | 0.032 |  |  |  |  |  |  |  |  |  |
| Dordogne | 0.059 | 0.055 | **0.013** | **0.008** |  |  |  |  |  |  |  |  |
| Garonne | 0.055 | 0.067 | **0.013** | **0.006** | **-0.001** |  |  |  |  |  |  |  |
| Loire | 0.046 | 0.052 | **0.032** | 0.019 | **0.006** | **0.009** |  |  |  |  |  |  |
| Vilaine | 0.073 | 0.056 | 0.029 | 0.032 | 0.018 | 0.022 | **0.005** |  |  |  |  |  |
| Scorff | 0.071 | 0.075 | 0.035 | 0.051 | 0.029 | 0.037 | 0.024 | 0.021 |  |  |  |  |
| Aulne | 0.101 | 0.109 | 0.056 | 0.049 | 0.062 | 0.052 | 0.047 | 0.052 | **0.023** |  |  |  |
| Trieux | 0.087 | 0.079 | 0.040 | 0.051 | 0.048 | 0.045 | 0.036 | 0.028 | **0.006** | **0.010** |  |  |
| Vire | 0.119 | 0.077 | **0.051** | 0.073 | 0.063 | 0.048 | 0.045 | 0.044 | 0.045 | 0.060 | 0.041 |  |
| Orne | 0.090 | 0.037 | **0.029** | **0.039** | **0.026** | **0.015** | **0.025** | 0.026 | 0.031 | **0.048** | **0.038** | **0.018** |

In *A. alosa* the highest likelihood (Ln(K)) was obtained for K=6 (**Fig. S2**), while ΔK showed a strong peak for K= 3 and another for K=6 (**Fig. S2**). Therefore we displayed the clustering results for both of these two values. For K=3 the clustering separated the populations into three geographic regions (Atlantic, Brittany, and Normandy/Nivelle, **Fig. 2A**). The Nivelle individuals (Southern France) clustered with individuals from Normandy in Northern France (Vire and Orne river in Fig. 2A) and this cluster was itself admixed with Brittany. The clusters from Brittany and Atlantic were themselves highly admixed between each other. For K=6 (**Fig. 2A**), the Nivelle individuals were separated from Normandy and formed a single cluster (contribution = 0.851). Normandy individuals formed a separated cluster, as well as Brittany individuals. Populations from the Atlantic were separated into three admixed clusters. Overall, most clusters were admixed with individuals from other genetic groups.

**Figure 2:**
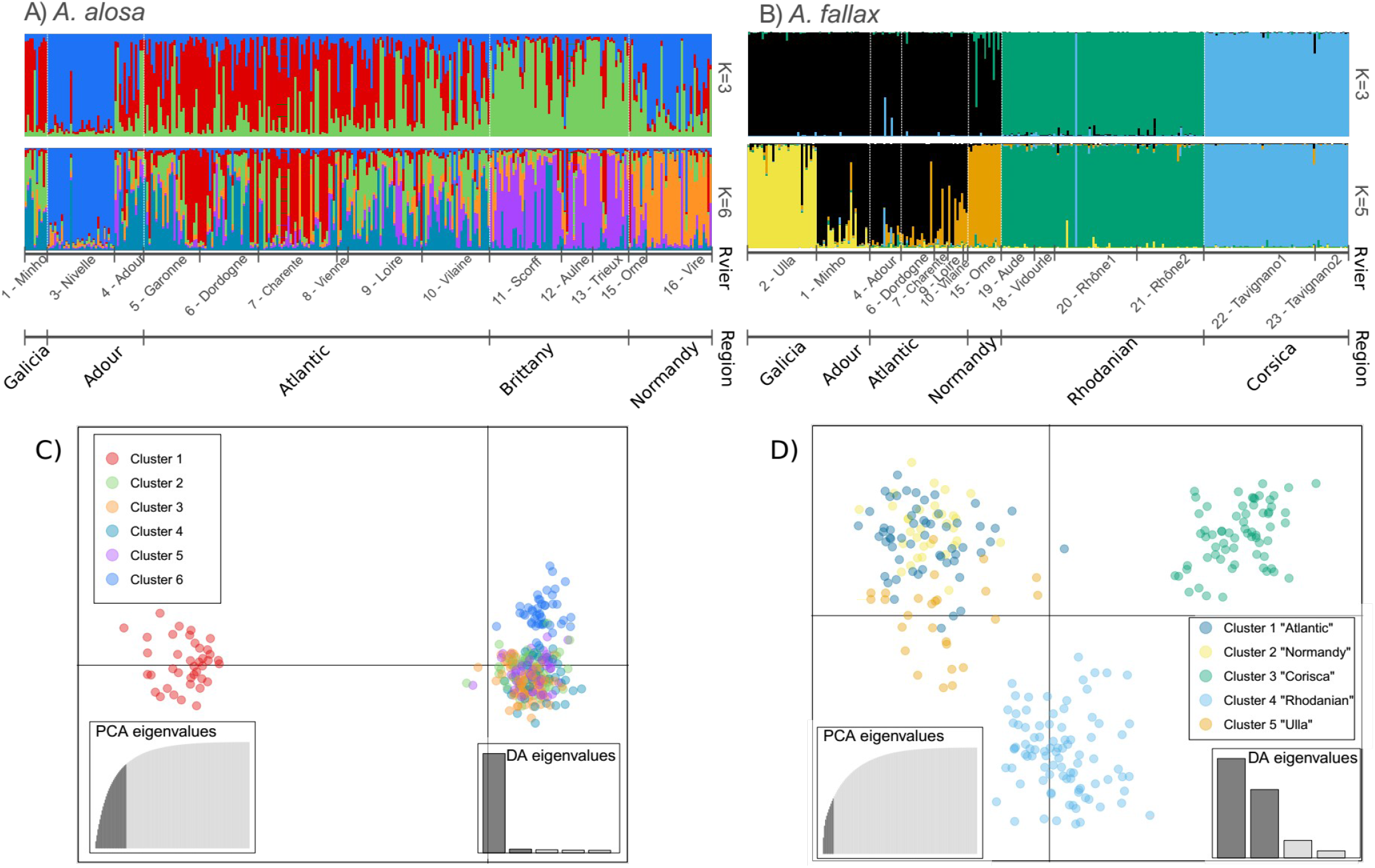
A) Bayesian individual clustering performed with STRUCTURE for each species separately. In A. alosa K = 3 and K=6 clusters are identified according to the L(K) and delta(K) methods presented in Fig S2. B) In A. fallax 3 and 6 clusters are identified according to the L(K) and delta(K) methods presented in Fig S2. Individuals are represented by a bar with each colour representing its membership proportion to each cluster. C) plot of the Discriminant Analysis of Principal Components (DAPC) performed in A. alosa. The “best” number of clusters was inferred using the Bayesian Information Criteria (BIC) (Fig S3). Clusters are numbered from one to six given that mixed membership probability was often attributed to different rivers. Membership assignment of individuals averaged by river to each cluster can be found in **Table S8**). D) plot of the DAPC performed in A. fallax. Membership assignment of individuals averaged by river to each cluster can be found in **Table S9**)

In *A. fallax*, the highest likelihood plateaued between K=5 and K=8, while ΔK showed a single strong peak for K=3 (**Fig. S3**). Therefore we present the results for K = 3 and K = 5 and we provide plots for K = 4 and K = 6 as supplementary results, although these were less supported. At K=3, individuals from the Atlantic, the North Mediterranean coast and Corsica were separated. In contrast to patterns observed in A. alosa, no sign of admixture was observed in *A. fallax* populations (**Fig. 2B**). Indeed, all individuals except those from the Orne River had >0.95 membership probability to a separate cluster (individuals in the Orne River still had membership probabilities to a distinct cluster greater than 0.86). With K=5, a separation of samples along the Atlantic Coast into 3 groups was revealed (Fig 2B). The Orne (Normandy) river formed one cluster and the Ulla (Galicia, Spain) formed a separate cluster. Increasing the number of groups to 6 separated the Minho from the remaining groups. Ts. The remaining populations from the French Atlantic Coast formed a single cluster on the Rhône River. One individual was assigned as a migrant from the Tavignano (individual q-value = 0.981 CI: 0.894 – 1.00). Although some individuals displayed admixed ancestry, only one individual from the Ulla River was assigned with probability greater than 0.8 to the Minho suggesting relatively low dispersal among clusters as opposed to the pattern observed in *A. alosa*. The DAPC approach revealed a similar level of population structure in both *A. alosa* (**Fig. 2C**) and *A. fallax* (**Fig 2D**). Indeed, according to the BIC, a total of 6 and 5 groups were present in each species respectively (Fig S4, Fig S5). However, the results for A. alosa were difficult to interpret (Table S9). Indeed, we find little congruence in the assignment of individuals to different groups in the DAPC as compared to structure. Only individuals from the Minho and from the Nivelle were assigned to a discrete cluster (Table S10) but these clusters displayed a very close relationship to other clusters in the DAPC plot (Fig 2).

To reveal potential hierarchical structure and fine scale genetic structure we replicated the analysis in *A. fallax* including only populations from the Atlantic coast. This revealed the existence of 4 clusters (**Fig. S3B**).

The neighbor-joining tree revealed a clear separation between *A. alosa* and *A. fallax* (**Fig. 3B**). In *A. alosa* 5 clusters can be delineated, with the Adour being separate and the Vilaine grouping with the Minho. In *A. fallax* three main clusters can be distinguished corresponding to a geographic clustering pattern. Separation of *A. fallax* from the Tavignano was as strong as the separation between the two cryptic lineages from the Mediterranean Sea and the Atlantic coast. Finally, tests for IBD were significant in both species (Mantel tests: P<0.001. r = 0.621 in *A. alosa* and P<0.0001. r = 0.554 in *A. fallax*. **Fig. 3A).** These results were also supported by significant linear models (**Fig 3A**).

**Figure 3:**
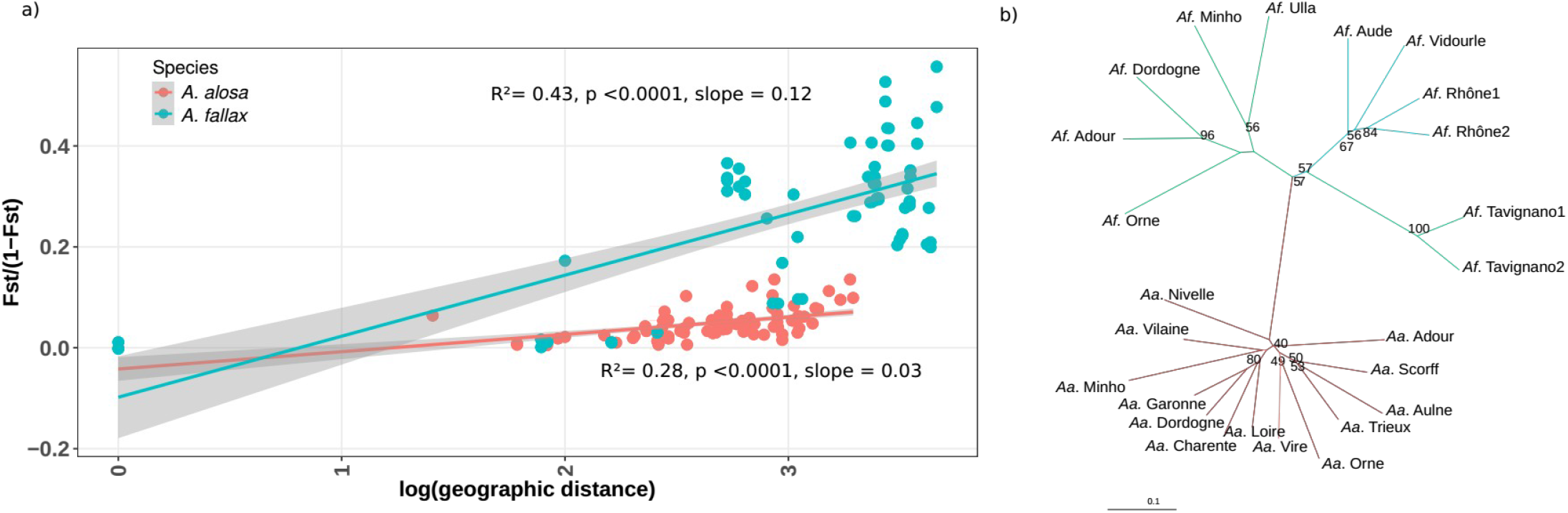
**A)** Relationship between genetic distance as measured by F_ST_/(1-F_ST_) and ln(distance) in kilometers between populations of each species. All correlations were significant based on linear models and Mantel tests. **B)** Neighbor-joining tree based on Cavalli-Svorza and Edwards (1967) genetic distances. Only bootstrap values >50% are indicated. Sample abbreviations match those given in Table 1. A**a** = A. alosa. A**f** = A. fallax.

### Assignment tests

According to GeneClass three individuals of *A. fallax* captured at sea displayed a probability <0.01 to originate from one of the sampled rivers and were excluded from further analyses with the three assignment methods. The forty *A. alosa* had a probability >0.01 to originate from one of the rivers. Contrasting results were obtained for each species; *A. fallax* were assigned with higher probability than *A. alosa* (only fish with score >0.9 were kept for assignment), with a total of 34 to 45 and of 18 to 34 fish respectively assigned depending on the method (**Table 3**). All methods assigned the majority of *A. alosa* to the Atlantic cluster. The DAPC approach classified 6 *A. alosa* from South Brittany in Normandy and 4 in the Minho, while these fish were either assigned to the Atlantic cluster or left unassigned by the other methods. Similarly, one *A. alosa* from the Pertuis Charentais was assigned to the Minho by DAPC while the two other methods classified this fish as belonging to the Atlantic cluster. In *A. fallax* the majority of fish from the North Sea, the South Brittany and the Southern part of the Bay of Biscay were assigned to the Orne cluster. Most fish from the Pertuis Charentais were assigned to the Atlantic cluster. One fish was assigned to the Ulla by the DAPC approach but was left unassigned by the two other methods. Similarly, two fish from Southern Brittany were assigned to the Minho with DAPC, whereas both were left unassigned by Structure and one was unassigned by GeneClass, while the other was assigned to the Atlantic cluster.

**Table 3:** Results of assignment test of A. alosa and A. fallax captured at sea. Shown are the proportions of individuals from a given sampling area assigned to a region (in the case of A. alosa) or to a river (A. fallax).

| Species | Sampling area | Number | Assignment Population | Structure | DAPC | GeneClass |
| --- | --- | --- | --- | --- | --- | --- |
|  |  |  |  | % | % | % |
| *A. Alosa* | South Brit-tany | 36 | Atlantic | 28 | 33 | 36 |
|  |  |  | Brittany | 11 | 8 | 11 |
|  |  |  | Normandy | 0 | 22 | 0 |
|  |  |  | Minho | 0 | 11 | 0 |
|  |  |  | Nivelle | 3 | 14 | 3 |
|  |  |  | Unassigned | 58 | 12 | 50 |
|  | Pertuis Charentais | 4 | Atlantic | 75 | 75 | 50 |
|  |  |  | Brittany | 0 | 0 | 25 |
|  |  |  | Minho | 0 | 25 | 0 |
|  |  |  | Normandy | 0 | 0 | 25 |
|  |  |  | Unassigned | 25 | 0 | 0 |
| *A. fallax* | South Brit-tany | 16 | Atlantic | 27 | 38 | 20 |
|  |  |  | Orne | 53 | 56 | 53 |
|  |  |  | Unassigned | 20 | 6 | 27 |
|  | Pertuis Charentais | 21 | Atlantic | 66 | 62 | 62 |
|  |  |  | Orne | 5 | 19 | 5 |
|  |  |  | Minho | 0 | 13 | 0 |
|  |  |  | Ulla | 0 | 5 | 0 |
|  |  |  | Unassigned | 29 | 1 | 33 |
|  | Bay of Biscay | 5 | Orne | 100 | 100 | 80 |
|  |  |  | Atlantic | 0 | 0 | 20 |
|  | North sea | 6 | Orne | 75 | 75 | 75 |
|  |  |  | Atlantic | 0 | 25 | 25 |
|  |  |  | Unassigned | 25 | 0 | 0 |
|  | North Brit-tany | 20 | Orne | 75 | 90 | 65 |
|  |  |  | Unassigned | 25 | 10 | 35 |

### Demographic history

Reconstruction of demographic history was performed i) between the two species along the *Atlantic* coast, as well as between the divergent lineages of *A. fallax*, namely between ii) *A. fallax* “Atlantic” and *A. fallax* “Mediterranean Sea” and between iii) A*. fallax* “Mediterranean Sea” and *A. fallax* “Corsica”. In all three cases, ABC model choice rejected SI and AM in favour of models with ongoing gene flow, with support for secondary contact (SC) in all cases (Table S11). For instance, in between species comparisons, posterior probabilities were *P_(SC)_* = 0.980 versus P(SI) =0.02; *P_(IM)_* = 0.851 versus P(SI) = 0.149 and. P_*(AM)*_ = 0.720 versus P(SI) = 0.20). For the sake of conciseness we only present results for between species comparisons, but the others are shown in Table S11. In each between species pairwise comparison, our cross-validation procedure revealed that the robustness was 1 for all comparisons (**Fig S6 A-F**). Comparison of the model with ancient gene flow against ongoing gene flow led to a rejection of the AM model (P_*(SC)*_ = 0.669 vs P(am) = 0.331; P_*(IM)*_ = 0.599 vs P(am) = 0.401) with a cross-validation procedure revealing a robustness of 1 (**Fig S6A, B**). Finally, comparing IM and SC leads to a higher posterior probability of the latter with P(SC) = 0.738 vs P(IM) = 0.262 and a robustness of 1 (**Fig 4a**, **Fig S7C,D**).

**Figure 4:**
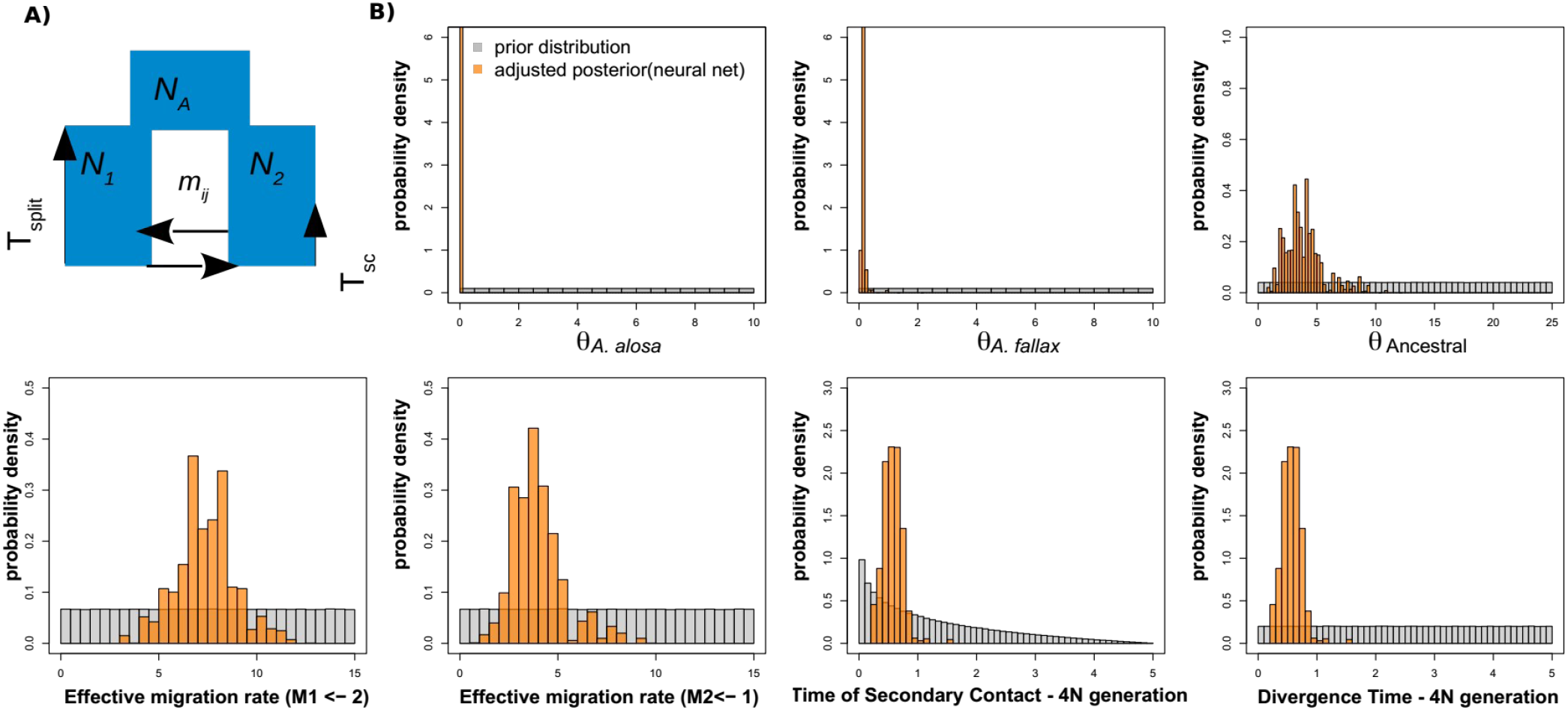
Schematic of the best model of divergence (panel A) and distributions of prior and posterior parameter estimates based on the model of best fit associated with divergence and diversity data (panel B). A) Simplified representation of the best inferred scenario of divergence between Alosa alosa and A. fallax. B) Prior (grey) and Posterior (orange) parameter distributions inferred by Approximate Bayesian Computation. θ A. alosa = Scaled Effective population size of A. Alosa. θ A. alosa = Scaled Effective population size of A. fallax. θ_ancestral_= Effective population size of the ancestral population. All populations size are scaled by 4Nref*μ. T_split_ = split time (here T_split_ = tau* 4N_ref_). T_sc_ = Time of Secondary Contact. M12 = 4Nref*m_12_ and M21 = 4Nref*m_21_ **corresponds** to the scaled migration rate where m_ij_ represents the fraction of subpopulation i that is made up of migrants from subpopulation j each generation. All values are provided in coalescent units and scaled by the effective reference population size (N_ref_ = 50,000).

The posterior distribution of parameter estimates associated with effective population size and divergence time under SC were well differentiated from the prior yielding confidence for interpretations of the mean values and credible intervals in between species comparisons (**Fig. 4b**) and between *A. fallax* lineages (Fig S8, Fig S9). These indicated population size reduction compared to the estimated ancestral effective population size (*N_ANC_*). The median estimated effective population size was *N_1_* = 209 [IC= 47 – 646] for *A. alosa, N_2_* = 2573 [940 – 5,649] for *A. fallax*, and *N_ANC_* = 617,000 [39,074 – 1,228,000] for the ancestral population (**Fig. 4b**). Credible intervals around the split time and the time of secondary contact overlapped. Median estimate of *T_split_* was ~774,000 generations but with large credible intervals [119,000 – 1.900,000]. Given the difference in life history, we did not convert these estimates into years. The time of secondary contact would be 157,000 generations [24,000 – 620,000] (Fig. 4b). Considering effective population size within *A. fallax*, we found that the Corsican lineage displayed the highest Ne (median = 19,500 [6,600 – 51,800]) and the Mediterranean lineage displayed a similar value to the Atlantic one (median = 3,000 [CI=100 – 9,600] depending on the comparison). Our inference further suggested that the divergence of *A. fallax* from the Atlantic and Mediterranean Sea was slightly more recent than that observed with *A. alosa* (median = 630,000 [CI = 69,000 – 970,000] generations) followed by a more recent divergence from *A. fallax* “Corsica” (median = 350,000 [CI = 35,000 – 960,000]. Additional parameter estimates such as migration rate can be found in Table S12 and Fig S8 and Fig S9.

## Discussion

Our results shed light on evolutionary processes affecting the rate of speciation and provide key information for the management of two declining fish species. We found that genetic structure among populations was lower for the nearly semelparous species, *A. alosa*, compared to the iteroparous species, *A. fallax*. Moreover, individuals captured at sea could be assigned at the region level for *A. alosa* and at the river level in *A. fallax*. We inferred that the most likely long term historical divergence scenario between both species implicated historical separation followed by a secondary contact accompanied by contemporary hybridization and strong population size declines. These observations regarding hybridization, demographic history and gene flow are in large agreement with those from Faria et al. (2012) which is the most recent broad scale study but uses mtDNA, thus bringing complementary insights into the species evolutionary dynamics. Our results also corroborate the hypothesis of a divergent lineage (*A. agone*) present along the Mediterranean coast and suggest a possible undocumented new cryptic lineage present in the Island of Corsica, which were both formerly grouped with *A. fallax*. These observations have strong conservation implications showing the importance of combining catchment and region-based management.

### Level of species differentiation and hybridization

Our results confirm previous estimates of introgression and hybridization between both species (Alexandrino et al. 2006; Coscia et al. 2010; Jolly et al. 2011; Faria et al. 2012; Taillebois et al. 2020). Normally, spatio-temporal segregation mechanisms exist and maintain or minimize the contact between the two species during reproduction. In particular, *A. fallax* generally spawns in lower areas of the watershed than *A. alosa*, which normally spawns in the upper parts of rivers (Aprahamian et al. 2003; Baglinière et al. 2003b), thereby minimizing hybridization opportunities between species. Yet, human river fragmentation by dams and various obstacles is increasingly restricting shad migration to downstream areas. This favors sympatry and leads to increased opportunities for hybridization. The removal of this premating reproductive barrier thus promotes hybridization and introgression (Alexandrino et al. 2006; Taillebois et al., 2020). Accordingly, we find strong support for ongoing hybridization between both species in populations distributed in the same watershed and reaching similar spawning sites (e.g. Loire and Charente River). The preferential occurrence of hybrids in these areas is unclear and may be related to the disruption of spawning grounds due to human activity, but this speculation would require further investigations. However, the two species seem to remain genetically distinct despite hybridization. This may indicate the existence of pre-zygotic (e.g. behavioral) and post-zygotic genetic barriers that are likely involved in the maintenance of reproductive isolation despite hybridization that seems to have existed over relatively long historical times (Ravinet et al. 2017, Barth et al. 2020). Yet, genetic barriers are often semi-permeable (Wu, 2001), resulting in heterogeneous differentiation across the genome (Ravinet et al. 2017; Rougemont et al. 2017). The porosity of the species barrier is well illustrated by the numerous backcrosses found recently by Taillebois et al. (2020) in a set of partially overlapping rivers using a SNP array as well as by the mtDNA results from Faria et al. (2012) on another set of overlapping rivers, thus providing support for our results. For instance, they identified bidirectional mitochondrial introgression ranging from 25 to 63% depending on the river considered (note that this percentage derived from a small number of individuals however). Whole genome re-sequencing of these hybridizing populations would be needed to i) quantify the mosaic of local ancestry across the genome (Duranton et al. 2018) ii) identify the regions involved in adaptive introgression from one species to the other (Hedrick 2013) and iii) determine the deleterious effect of introgression and extent of selection against introgression (Kim et al. 2018). The hybridization of these two species is hence an interesting context to study speciation and also highlights the consequences of river fragmentation on the genetic integrity of these declining species (Harris et al. 2018; Rougemont et al., 2020).

### Influence of life history strategy on the population genetic structure

As expected from theory and from previous research, the species that disperses more readily (and semelparous), *A. alosa*, displays lower levels of genetic structure and higher levels of genetic diversity compared to *A. fallax*. A limit here is that the inference of the number of clusters in *A. alosa* was difficult with structure analysis pointing to either three or six clusters.Therefore it is likely that the number of real ancestral groups for *A. alosa* is below the 3 or 6 groups inferred here. Another complicating factor was the significant signal of isolation by distance which itself can generate genetic clusters (Meirmans, 2012; Battey et al. 2020). Regardless of the exact number of clusters, *A. alosa* is likely to derive from one major group with signals of admixture being inflated due to IBD. These variations in dispersal syndrome are well documented and are associated with variation in life history traits across many species (Stevens et al. 2013; Cayuela et al. 2016, 2018). Here, this suggests a substantial trade-off associated with these life history traits. Since *A. fallaxis* iteroparous, we hypothesized that they maximize survival probability by dispersing less and reducing the duration of their marine phase due to an earlier age of sexual maturity than *A. alosa* (Bagliniere et al. 2020). Other hypotheses have been made to explain the weaker genetic structure of the semelparous American shad (*A. sapidissima*) as compared to its iteroparous form and relate to environmental differences along a latitudinal gradient (discussed in Hasselman et al. 2013). Here, however, both species reproduce in the same rivers along the Atlantic coast, so the proposed environmental mechanisms are less likely to apply in our case. A meta-analysis in terrestrial and semi-terrestrial animals revealed that higher dispersal was associated with higher fecundity and survival (Stevens et al. 2014). In a less stable environment, higher dispersal, higher fecundity and shorter lifespan can be favored (Cayuela et al. 2016). This leads us to hypothesize that *A. alosa* populations may be more resilient under climate change, but this hypothesis remains to be investigated.

### Demography and speciation

The differentiation levels among *A. fallax* lineages are similar to those observed between species. This led us to identify two putative cryptic species in addition to the already documented *A. fallax* located along the Atlantic coast. The first putative lineage occurs in the continental coast of the Mediterranean and the second one in Corsica. One caveat stems from our limited sampling of *A. fallax* between the Vidourle and Southern Spain. With such sparse sampling, genetic clustering methods can be confounded by isolation by distance as well as coalescent based species delimitation methods (Meirmans, 2012, Mason et al. 2020). Yet, our results are also supported by the observed increased differentiation, as well as the genetic differences between Atlantic and Mediterranean *A. fallax* based on protein markers and morphological differences observed by Le Corre et al. (2005). They also suggested that the Mediterranean *A. fallax* may be considered as an independent species, *Alosa agone* (Scopoli 1786). Indeed, based on the genetic studies of Le Corre et al. (2015) and morphological observations of Bianco et al. (2005), Mediterranean *A. fallax* was considered as a separate species (*A. agone*) in the recent atlas of freshwater fish from France. Overall the taxonomic status of this species is not yet firmly established (Chiesa et al. 2014; Baglinière et al. 2020c). While it is considered a valid species in the atlas of fish, previous genetic studies failed to distinguish *A. agone* from *A. fallax* (Faria et al. 2006; Chiesa et al. 2014) and our study is the first to provide evidence for strong differentiation between *A. fallax* and *A. agone.”* Additional examination of phenotypic and ecological differences between this lineage and the other A. fallax lineages, along with sampling of remote lineages (e.g. *Alosa immaculata*) would enable a better description of this species. The identification of a putative lineage in Corsica is a new finding. A parallel can be drawn with *Salmo trutta*, which is also represented with many phylogeographic lineages (Bernatchez 2001) and locally differentiated populations in the Tyrrhenian area, including Corsica, that are attached to the Adriatic lineage rather than the Mediterranean lineage (Berrebi et al. 2019). Unfortunately, microsatellite data are poorly suited to disentangle recently high genetic drift from long divergence time. A whole genome approach would be necessary to better reconstruct the divergence history of all the species and sub-populations or cryptic lineages using explicit modeling and additional measures such as absolute divergence (D_XY_; Nei, 1987). Presently, the systematics of the genus *Alosa* remain unclear because of the large number of subspecies of *A. fallax* (Chiesa et al. 2014; Coscia et al. 2013; Baglinière et al. 2020c).

In addition, we have attempted to reconstruct the demographic history of the two species and the divergent *A. fallax* lineages, though Yet we call attention to a few caveats. First, we had too few markers to highlight the semi-permeable nature of the genome and to discriminate local regions of the genome that are freely exchanged from those that are possibly impermeable to gene flow and may be involved in reproductive isolation (Duranton et al. 2018). A. alosa Moreover, the mutation rate is not known and the generation time remains difficult to estimate given that *A. alosa* reproduces only once between 3 to 8 years, whereas *A. fallax* can reproduce multiple times between 2 to 8 years (Baglinière et al. 2020). Assuming the same generation time as Faria et al. (2012), namely five years, we would obtain a divergence time of ~3My [95%CI: 500 Ky – 4.6My], thus compatible with estimates [0.294 – 3.477]My obtained in their study. Regardless of the timing of divergence, Faria et al. (2012) net sequence divergence (Da = 0.02) can be used to replace the species along the speciation continuum proposed by Roux et al. (2016). Their results thus indicate that the species fall exactly in the gray zone of speciation where reconstructing the divergence history is likely to be most difficult. Interestingly, the estimated time of divergence among species and lineages differ only by a small magnitude and considering the uncertainty associated with these parameters it is possible that all species and lineages have diverged simultaneously. More insights about the full species radiation history would require sampling all possible lineages within the Mediterranean and Black sea along with whole genome sequencing. Moreover, our estimates of the difference in effective population size between *A. fallax* and *A. alosa* as well as when compared to the ancestral population are also in broad agreement with those of Faria et al. (2012), although the exact value differs. This discrepancy may be due to the difference in maker resolution, difference in selective constraints undergone by mtDNA versus microsatellite markers or the difference in the methods used for demographic reconstruction. For instance, Faria et al. (2012) used IMa2, which is sensitive to confounding by linked selection (Cruickshank et Hahn, 2014). Most importantly, we used individuals from a single river to avoid increasing local population structure, whereas Faria et al. (2012) included samples from several rivers, and thus estimated the metapopulation Ne, preventing any direct comparison with our estimates.

Among other limits, our sampling design did not include *A. immaculata*, a species from the Caspian and Black sea that is more closely related to *A. fallax* than to *A. alosa* according to mtDNA analyses (Faria et al. 2012). However, the node support for their grouping was weak and did not entirely allow us to conclude that *A. fallax* originate from the same area as *A. immaculata* (Alexandrino et al. 2006; Faria et al. 2006; Faria et al. 2012). Therefore, the hypothesis of a shared origin in the Mediterranean Sea for *A. fallax* and *A. immaculata* versus an Atlantic origin for *A. alosa* cannot be excluded. This would lend support to the secondary contact model that we inferred here. Moreover, the absence of *A. immaculata* in our samples (i.e. ghost species) can influence our inferences of gene-flow (Mason et al. 2020; Tricou et al. 2022). For instance, ancient gene flow between *A. immaculata* and *A. alosa* may have left some footprint in the genome of *A. alosa* that may complicate parameter estimation. Yet, *A. alosa* and *A. fallax* are also known to share spawning grounds (Maitland et Lyle, 2005) and produce hybrids (e.g. Taillebois et al. 2020). Therefore, we also expect ongoing gene flow due to secondary contact to be a supported scenario. For instance, a chromosome painting approach (Lawson et al. 2012) using whole genome sequence data with all potential donor populations may help identify the origin of different ancestry tracks along the genome of *A. alosa*.

With all these caveats in mind, the higher support for a model of secondary contact between the two species than for alternative models, indicates that isolation was necessary to initiate divergence between the two species. Most theories suggest that speciation is difficult to establish in the presence of continuous gene flow (Barton and Bengtsson 1986; Bierne et al. 2011). Only a few studies convincingly support speciation with gene flow and have explicitly attempted to discriminate among competing models (Martin et al. 2013; Malinsky et al. 2015; Tusso et al. 2021). In contrast, an increasing number of studies have reported evidence for divergence initiated in allopatry followed by gene flow (e.g. Roux et al. 2013, 2014; Rougemont et al. 2017; Rougemont and Bernatchez 2018; Leroy et al. 2019; Cayuela et al. 2020). Interestingly, Tine et al. (2014) and Duranton et al. (2018) have reported the existence of two cryptic species of sea bass (*Dicentrarchus labrax*), the Atlantic and Mediterranean Sea Bass with evidence of secondary contact (Tine et al. 2014), multiple islands of reproductive isolation (Duranton et al. 2018), and cryptic genetic and demographic connectivity (Robinet et al. 2020). Similarly, Riquet et al. (2019) reported the existence of divergent lineages of seahorses (*Hippocampus guttulatus*) on the Mediterranean Sea and the Atlantic coast as well as the existence of partially reproductively isolated cryptic lineages maintained in sympatry within the Mediterranean Sea. Shared origin in the mediterranean sea for *A. fallax* and *A. immaculata* versus an Atlantic origin for *A. alosa* cannot be excluded. This would lend support to the secondary contact model that we inferred here. Moreover, the absence of *A. immaculata* in our samples (i.e. ghost species) can influence our inferences of gene-flow (Mason et al. 2020; Tricou et al. 2022). For instance, ancient gene flow between *A. immaculata* and *A. alosa* may have left some footprint in the genome of *A. alosa* that may complicate parameter estimation. Yet, *A. alosa* and *A. fallax* are also known to share spawning ground and to produce hybrids (e.g. Taillebois et al. 2019). Therefore, we also expect ongoing gene flow due to secondary contact to be a supported scenario. For instance, a chromosome painting approach (Falush et al. 2012) using whole genome sequence data with all potential donor populations may help identify the origin of different ancestry tracks along the genome of *A. alosa*. Finally, our results have important conservation implications. First, our estimates of long-term effective population size indicated that the *Ne* of *A. fallax* was higher than that of *A. alosa*. This estimate is a long-term average based on the number of coalescent events that have occurred until all lineages coalesce. It reflects the historical population size and is in line with previous phylogeographic knowledge (Alexandrino et al. 2006; Faria et al. 2012). Despite all the uncertainty associated with ancestral population size estimates, our results indicate that both *A. fallax* and *A. alosa* have undergone strong reductions following their divergence. Therefore, the ongoing decline of the two species (Aprahamian et al. 2003; Baglinière et al., 2003b; Rougier et al., 2012) is of further concern if they already have a reduced adaptive potential. These long-term trends further suggest a reduced evolutionary potential (Leroy et al. 2021). Finally, we were able to identify and assign individuals captured at sea of unknown origin to putative river or regional groups with modest power. These results suggest that our marker set could be improved (e.g. using additional markers or by moving to a SNP array) to identify individuals dispersing at sea and draw inferences about dispersal distances of the two species, an important problem when assessing the provenance of individuals in mixed-stocks fisheries (Beacham et al. 2019; Nachón et al. 2020).

### Conclusion and future directions

Our results indicate that *Alosa* species constitute an interesting model to study speciation and hybridization between closely related species. Understanding the fitness effects of such ongoing hybridization will be of utmost importance to help manage the species. For instance, if the species are increasingly constrained to breed on overlapping spawning sites, the rate of hybridization is expected to increase, which may lead to increased fitness of the hybrid during the first few generations of hybridization, but may also increase selection against introgression if recessive deleterious alleles are expressed. This could negatively affect population size, by reducing the fitness of backcrosses and hence decrease the species’ evolutionary potential. However, the application of the EU Water Framework Directive on good ecological status could reverse the hybridization trend of these species if the ecological connectivity of rivers is restored and the natural, separate breeding grounds of both species are accessible again. Overall, our results revealed deep divergence followed by secondary contact and a prevailing role of gene flow between the two species as well as among new cryptic lineages or possible species of *A. fallax*. These results highlighted the consequences of contrasting life history strategies at the genetic level and suggest that the two species should be managed jointly given their porous reproductive boundaries. We propose that whole genome sequences will help address several questions that have been raised, especially the inference of historical divergence and demography of these species, and the inference of putative post-zygotic barriers across the genome.

## Acknowledgments

We thank the many professional fishermen involved in gathering samples. We thank the institutions involved in collecting samples, namely people at INRAE, OFB and FDPPMA 14, 22, 50, and Migado association. We thank A. Xuereb for extensive revision of the manuscript grammar. This study was funded by the European Regional Development Fund (Transnational program Interreg IV. Atlantic Aquatic Resource Conservation Project).

## Data accessibility

data will be deposited on dryad and code for ABC simulations are available on github.

