## supplementary material for "Population genetics reveals divergent lineages and ongoing hybridization in a declining migratory fish species complex"

### Supplementary Figures and Tables

**Supplementary Table S1:** Details of sample sites and samples collected at sea

**Supplementary Table S2:** Genetic diversity observed in *A. alosa* at each microsatellite marker (1 column = 1 marker). Details are displayed for each river (rows) and contains the following information: N = Number of individuals Sampled, A = Number of Alleles, Ar = Allelic Richness, Ho = Observed Heterozygosity, He = Expected Heterozygosity, Fis = Inbreeding coefficient, HWE = p-value of being out of HWE.

**Supplementary Table S3:** Genetic diversity observed in *A. fallax* at each microsatellite marker (1 column = 1 marker). Details are displayed for each river (rows) and contains the following information: N = Number of individuals Sampled, A = Number of Alleles, Ar = Allelic Richness, Ho = Observed Heterozygosity, He = Expected Heterozygosity, Fis = Inbreeding coefficient, HWE = p-value of being out of HWE.

**Supplementary Table S4:** Results of the mixed linear models (LMM) testing for difference in allelic diversity (response variable = Allelic Richness) among populations.

**Supplementary Table S5:** Results of Tukey HSD test for differences in Allelic Richness among populations.

**Supplementary Table S6:** Mean Efficiency and Accuracy of hybrid identification for Structure and NewHybrid when considering different q-value threshold obtained on simulated data from HybridLab.

**Supplementary Table S7:** q-values of individuals classified as hybrids according to *New Hybrids* as well as q-value from Structure along with their 90% confidence intervals (available only in Structure). Three categories were considered in *New hybrids* corresponding either to i) pure *A. alosa*, ii) pure *A. fallax* or iii) *hybrid*. Hybrids from new *NewHybrids* could correspond to either F1, F2 or backcrosses and were lumped into a single category given that the highest power and efficiency were reached when these categories were merged. In Structure two categories were possible, corresponding to either *A. fallax* or *A. alosa*, with hybrid expect to displayed intermediate q-values (from 0.25 to 0.75 depending on the directions of introgression).

**Supplementary Table S8:** Pairwise weir & cockerham $F_{ST}$ between all populations and between species.

**Supplementary Table S9:** Mean membership assignment of *A. alosa* individuals averaged by river to each cluster found in the DAPC displayed in Fig 2c.

**Supplementary Table S10:** Mean membership assignment of *A. fallax* individuals averaged by river to each cluster found in the DAPC displayed in Fig 2d.

**Supplementary Table S11: Model choice for each species and lineages.** Posterior probabilities (P) of each model and its alternative were obtained through the neural network approach implemented in the ABC package. Values are display for the between-species comparison as well as between lineages of *A. fallax*. At the first round of comparison model with gene flow were compared against the model of strict isolation. At the second round, then the two model with ongoing gene-flow (IM and SC) were compared against model with Ancient gene-flow (AM). Finally, the two best model (IM and SC) were compared against each other at the last round. AM = Ancient Migration, IM = Isolation with Migration, SC = Secondary Contact SI = Strict Isolation.

**Supplementary Table S12:** Prior and posterior parameter estimates for ABC computations. Uniform prior (U) distribution were used. The values displayed for the rate of migration are scale by $4 * N_{ref} * \mu$ assuming $N_{ref} = 50,000$ and $\mu = 2.5e-4$ mutations/bp/generation, as coalescent simulation are always scaled in the coalescent simulator used (ms).

$N_{allis\_shad}$ = effective population size of Allis shad (*A. alosa*). $N_{rwaite\_shad}$ = effective population size of twaite shad (*A. fallax*) $m1$ = effective migration rate from population 2 into population 1, $m2$ = effective migration rate population 1 into population 2. Here population 1 = *A. alosa* and population 2 = *A. fallax*.

$T_{split}$ = Divergence Time between species (in generations) and $T_{sc}$ = Time of secondary contact between the two species.

**Supplementary Figure S1:** Compared demographic models of divergence

**Figure S2: Results of Structure model choice.** A) Delta K for *A. alosa* obtained using Evanno et al. (2005) B) L(K) for *A. alosa* obtained using pritchard et al. (1999) method, C) Delta K for *A. fallax* and D) L(K) for *A. fallax*

**Supplementary Figure S3:** A) Ln (K) and B) delta K obtained when considering *A. fallax* sampled along the Atlantic coast only and C) corresponding population genetic structure plot.

**Supplementary Figure S4: Bayesian Information Criteria (BIC) revealing the number of group used in the DAPC and α-score obtained in A) *A. alosa* and B) *A. fallax*.**

**Supplementary Figure S5: Robustness of the model choice when comparing isolation model (SI) against models with ongoing gene flow (IM, panel A,B) ; Ancient Migration (AM, panel C,D) or Secondary Contact (SC, panel D,E).** Robustness is obtained through 4,000 cross-validations using the ABC model choice procedure (neural networks) on simulated data.

A) Posterior probability distribution densities of inferring the true model (M) given that the model was generated under M, i.e. $P(M|M)$. M being either AM (yellow line) or IM (black line)

B) Posterior probability of observing the IM model given it was generated under either IM or AM.

C) Posterior probability distribution densities of inferring the true model (M) given that the model was generated under M, i.e. $P(M|M)$. M being either IM (yellow line) or SC (black line)

D) Posterior probability of observing the SC model given it was generated under either IM or SC.

Red line = posterior probability of the model observed in the empirical data.

**Supplementary Figure S6: Robustness of the model choice comparing model with ongoing gene-flow (IM) against models with Ancient Gene flow (AM) or secondary Contact (SC).**

Robustness is obtained through 4,000 cross-validations using the ABC model choice procedure (neural networks) on simulated data.

A) Posterior probability distribution densities of inferring the true model (M) given that the model was generated under M, i.e. $P(M|M)$. M being either AM (yellow line) or IM (black line)

B) Posterior probability of observing the IM model given it was generated under either IM or AM.

C) Posterior probability distribution densities of inferring the true model (M) given that the model was generated under M, i.e. $P(M|M)$. M being either IM (yellow line) or SC (black line)

D) Posterior probability of observing the SC model given it was generated under either IM or SC.

Red line = posterior probability of the model observed in the empirical data.

**Supplementary Figure S7: Prior (grey) and Posterior (orange) distribution of parameter estimates associated to the best model inferred by ABC when considering *A. fallax* lineages:**

$\theta A. fallax$ Atlantic = Scaled Effective population size of *A. fallax* "Atlantic lineage". $\theta A. fallax$ "Mediterranean" = Scaled Effective population size of *A. fallax* "Mediterranean lineage". $\theta_{ancestral}$ = Effective population size of the ancestral population. All populations size are scaled by $4N_{ref}\mu$. $T_{split}$ = split time (here $T_{split} = \tau * 4N_{ref}$). $T_{sc}$ = Time of Secondary Contact. $M_{12} = 4N_{ref}m_{12}$ and $M_{21} = 4N_{ref}m_{21}$ correspond to the scaled migration rate where $m_{ij}$ represents the fraction of subpopulation $i$ which is made up of migrants from subpopulation $j$ each generation. All values are provided in coalescent units and scaled by the effective reference population size ($N_{ref} = 50,000$).

**Figure S8: Prior (grey) and Posterior (orange) distribution of parameter estimates associated to the best model inferred by ABC when considering *A. fallax* lineages within Mediterranean Sea:**

$\theta A. fallax$ "Mediterranean" = Scaled Effective population size of *A. fallax* "Mediterranean lineage". $\theta A. fallax$ "Corsica" = Scaled Effective population size of *A. fallax* "Corsican lineage". $\theta_{ancestral}$ = Effective population size of the ancestral population. All populations size are scaled by $4N_{ref}\mu$. $T_{split}$ = split time (here $T_{split} = \tau * 4N_{ref}$). $T_{sc}$ = Time of Secondary Contact. $M_{12} = 4N_{ref}m_{12}$ and $M_{21} = 4N_{ref}m_{21}$ correspond to the scaled migration rate where $m_{ij}$ represents the fraction of subpopulation $i$ which is made up of migrants from subpopulation $j$ each generation. All values are provided in coalescent units and scaled by the effective reference population size ($N_{ref} = 50,000$).

**Supplementary Table S1:** Details of samples collected at sea

N = Number of individuals. The sample locations numbers (Sample ID) are also displayed on Figure 1.

|  | **Individuals at sea:** |  |  |  |
| --- | --- | --- | --- | --- |
| ***Species*** | **Sampling area** | **Sample ID** | **N.** | **N. without hybrid** |
| ***A. alosa*** | South Brittany | 26 | 36 | 36 |
|  | Pertuis Charentais | 25 | 4 | 4 |
| ***A. fallax*** | South Brittany | 26 | 16 | 15 |
|  | Pertuis Charentais | 25 | 21 | 21 |
|  | Bay of Biscay | 24 | 5 | 5 |
|  | North Brittany | 27 | 20 | 20 |
|  | North Sea | 28 | 6 | 4 |
|  | ***total A. alosa*** |  | 40 | 40 |
|  | ***total A. fallax*** |  | 68 | 65 |

**Supplementary Table S2:** Genetic diversity observed in *A. alosa* at each microsatellite markers (1 column = 1 markers). Details are displayed for each river (rows) and contains the following information: N = Number of individuals Sampled, A = Number of Alleles, Ar = Allelic Richness, Ho = Observed Heterozygosity, He = Expected Heterozygosity, Fis = Inbreeding coefficient, HWE = pvalue of being out of HWE. River Number match those on the map of Figure 1.

| Region | River + ID |  | Alo<br>01 | Alo<br>06 | Alo<br>07 | Alo0<br>9 | Alo15 | Alo1<br>6 | Alo<br>17 | Alo<br>26 | Alo<br>29 | Alo<br>32 | Alo3<br>3 | Alo4<br>3 | Alo45 |
| --- | --- | --- | --- | --- | --- | --- | --- | --- | --- | --- | --- | --- | --- | --- | --- |
| Spain | 1 - Minho | N | 10 | 10 | 10 | 10 | 10 | 10 | 10 | 10 | 10 | 10 | 10 | 10 | 10 |
|  |  | A | 7 | 6 | 6 | 7 | 4 | 2 | 3 | 5 | 6 | 4 | 7 | 5 | 5 |
|  |  | Ar | 6.77 | 4.73 | 5.4 | 6.33 | 3.77 | 1.97 | 2.8 | 4.57 | 4.6 | 3.97 | 5.59 | 4.77 | 4.8 |
|  |  | Ho | 0.56 | 0.6 | 0.9 | 0.7 | 0.5 | 0.2 | 0.4 | 0.5 | 0.9 | 0.6 | 0.9 | 0.9 | 0.9 |
|  |  | He | 0.86 | 0.63 | 0.76 | 0.74 | 0.55 | 0.19 | 0.35 | 0.65 | 0.75 | 0.71 | 0.79 | 0.76 | 0.77 |
|  |  | Fis | 0.49 | 0.13 | -0.31 | 0.07 | 0.17 | 0.42 | -0.14 | 0.35 | -0.12 | 0.16 | -0.12 | -0.16 | -0.17 |
|  |  | HWE | 0.01 | 0.12 | 0.47 | 0.54 | 0.34 | 1 | 1 | 0.21 | 0.96 | 0.18 | 0.94 | 0.5 | 0.94 |
| southern<br>France | 3- Nivelle | N | 30 | 30 | 30 | 30 | 30 | 30 | 30 | 30 | 30 | 30 | 30 | 30 | 30 |
|  |  | A | 7 | 4 | 4 | 10 | 7 | 2 | 2 | 5 | 4 | 6 | 6 | 7 | 5 |
|  |  | Ar | 4.56 | 3.84 | 3.41 | 7.12 | 3.61 | 2 | 2 | 3.52 | 2.53 | 5.21 | 3.97 | 5.42 | 4.11 |
|  |  | Ho | 0.67 | 0.67 | 0.48 | 0.8 | 0.7 | 0.43 | 0.53 | 0.47 | 0.27 | 0.77 | 0.67 | 0.8 | 0.67 |
|  |  | He | 0.73 | 0.71 | 0.45 | 0.87 | 0.61 | 0.41 | 0.51 | 0.58 | 0.24 | 0.8 | 0.56 | 0.8 | 0.72 |
|  |  | Fis | 0.09 | 0.06 | 0.06 | 0.08 | -0.12 | 0.02 | -0.05 | 0.21 | -0.09 | 0 | -0.22 | -0.01 | 0.04 |
|  |  | HWE | 0.43 | 0.51 | 0.79 | 0.04 | 0.87 | 0.76 | 0.74 | 0.26 | 1 | 0.44 | 0.98 | 0.37 | 0.13 |
|  | 4 - Adour | N | 13 | 13 | 13 | 13 | 13 | 13 | 13 | 13 | 13 | 13 | 13 | 13 | 13 |
|  |  | A | 7 | 4 | 3 | 7 | 3 | 2 | 3 | 5 | 6 | 5 | 6 | 5 | 7 |
|  |  | Ar | 5.65 | 3.23 | 2.57 | 5.89 | 2.62 | 1.99 | 2.62 | 4.56 | 5.2 | 4.46 | 5.7 | 4.7 | 6.09 |
|  |  | Ho | 0.69 | 0.54 | 0.31 | 0.67 | 0.69 | 0.31 | 0.54 | 0.54 | 0.75 | 0.54 | 0.7 | 0.75 | 0.91 |
|  |  | He | 0.69 | 0.58 | 0.28 | 0.8 | 0.55 | 0.27 | 0.5 | 0.74 | 0.75 | 0.71 | 0.73 | 0.72 | 0.73 |
|  |  | Fis | -0.01 | 0.08 | -0.1 | 0.17 | -0.27 | -0.14 | -0.09 | 0.28 | -0.01 | 0.25 | 0.05 | -0.04 | -0.26 |
|  |  | HWE | 0.67 | 0.49 | 1 | 0.21 | 0.9 | 1 | 0.79 | 0.02 | 0.43 | 0.19 | 0.27 | 0.7 | 1 |
| Atlantic | 5 - Garonne | N | 25 | 25 | 25 | 25 | 25 | 25 | 25 | 25 | 25 | 25 | 25 | 25 | 25 |
|  |  | A | 12 | 7 | 6 | 9 | 5 | 2 | 2 | 5 | 10 | 4 | 9 | 7 | 7 |
|  |  | Ar | 7.7 | 3.65 | 3.76 | 6.05 | 3.34 | 2 | 1.98 | 3.99 | 6.61 | 3.49 | 6.49 | 5.6 | 5.34 |
|  |  | Ho | 0.92 | 0.44 | 0.5 | 0.72 | 0.6 | 0.4 | 0.28 | 0.64 | 0.6 | 0.72 | 0.72 | 0.8 | 0.76 |
|  |  | He | 0.87 | 0.6 | 0.48 | 0.74 | 0.59 | 0.37 | 0.3 | 0.67 | 0.79 | 0.56 | 0.77 | 0.78 | 0.8 |
|  |  | Fis | -0.05 | 0.3 | -0.03 | -0.01 | -0.01 | -0.08 | 0.13 | 0.07 | 0.2 | -0.33 | 0.05 | 0.01 | 0.04 |
|  |  | HWE | 0.54 | 0.09 | 0.75 | 0.48 | 0.63 | 0.82 | 0.58 | 0.15 | 0.05 | 1 | 0.4 | 0.73 | 0.29 |
|  | 6 - Dordogne | N | 26 | 26 | 26 | 26 | 26 | 26 | 26 | 26 | 26 | 26 | 26 | 26 | 26 |
|  |  | A | 11 | 7 | 8 | 10 | 5 | 3 | 2 | 4 | 12 | 5 | 7 | 7 | 6 |
|  |  | Ar | 7.69 | 5.05 | 4.59 | 6.51 | 3.29 | 2.35 | 2 | 3.58 | 7.76 | 3.65 | 5.48 | 4.92 | 4.64 |
|  |  | Ho | 0.88 | 0.69 | 0.52 | 0.81 | 0.5 | 0.43 | 0.38 | 0.65 | 0.72 | 0.62 | 0.68 | 0.63 | 0.58 |
|  |  | He | 0.87 | 0.74 | 0.55 | 0.82 | 0.57 | 0.48 | 0.43 | 0.65 | 0.85 | 0.53 | 0.73 | 0.71 | 0.71 |
|  |  | Fis | -0.02 | 0.06 | 0.14 | 0.02 | 0.12 | 0.16 | 0.17 | 0 | 0.14 | -0.13 | 0.1 | 0.07 | 0.19 |
|  |  | HWE | 0.73 | 0.47 | 0.18 | 0.54 | 0.25 | 0.43 | 0.44 | 0.63 | 0.07 | 0.77 | 0.09 | 0.04 | 0.08 |
|  | 7 - Charente | N | 35 | 35 | 35 | 35 | 35 | 35 | 35 | 35 | 35 | 35 | 35 | 35 | 35 |
|  |  | A | 15 | 9 | 7 | 8 | 7 | 3 | 4 | 5 | 11 | 7 | 9 | 9 | 6 |
|  |  | Ar | 6.89 | 5.51 | 3.64 | 5.32 | 3.52 | 2.42 | 2.23 | 4.17 | 6.32 | 4.1 | 5.49 | 5.14 | 5.25 |
|  |  | Ho | 0.83 | 0.63 | 0.35 | 0.73 | 0.54 | 0.35 | 0.29 | 0.51 | 0.64 | 0.57 | 0.63 | 0.66 | 0.71 |
|  |  | He | 0.82 | 0.78 | 0.38 | 0.71 | 0.59 | 0.44 | 0.39 | 0.63 | 0.83 | 0.57 | 0.73 | 0.69 | 0.79 |
|  |  | Fis | 0.02 | 0.25 | 0 | -0.01 | 0.13 | 0.19 | 0.14 | 0.19 | 0.21 | 0.07 | 0.13 | 0.07 | 0.09 |
|  |  | HWE | 0.41 | 0 | 0.02 | 0.73 | 0.38 | 0.15 | 0.09 | 0.04 | 0.02 | 0.47 | 0.17 | 0.26 | 0.14 |
|  | 9 - Loire | N | 33 | 33 | 33 | 33 | 33 | 33 | 33 | 33 | 33 | 33 | 33 | 33 | 33 |
|  |  | A | 12 | 7 | 9 | 10 | 7 | 4 | 4 | 5 | 8 | 5 | 8 | 10 | 8 |
|  |  | Ar | 6.5 | 4.91 | 4.27 | 6.7 | 4.12 | 2.67 | 2.46 | 4.05 | 4.28 | 3.89 | 5.84 | 6.15 | 5.78 |
|  |  | Ho | 0.73 | 0.7 | 0.45 | 0.75 | 0.64 | 0.55 | 0.39 | 0.63 | 0.33 | 0.52 | 0.74 | 0.67 | 0.85 |
|  |  | He | 0.81 | 0.72 | 0.46 | 0.81 | 0.68 | 0.54 | 0.35 | 0.66 | 0.54 | 0.59 | 0.79 | 0.72 | 0.78 |
|  |  | Fis | 0.07 | 0.02 | 0 | 0.07 | 0.05 | -0.06 | -0.15 | 0.02 | 0.39 | 0.15 | 0.07 | 0.09 | -0.08 |

|  |  | HWE | 0.04 | 0.5 | 0.33 | 0.05 | 0.01 | 0.59 | 0.83 | 0.01 | 0 | 0.26 | 0.38 | 0.18 | 0.91 |  |
| --- | --- | --- | --- | --- | --- | --- | --- | --- | --- | --- | --- | --- | --- | --- | --- | --- |
|  | 10 - Vilaine | N | 30 | 30 | 30 | 30 | 30 | 30 | 30 | 30 | 30 | 30 | 30 | 30 | 30 |  |
| Brittany |  | A | 12 | 5 | 5 | 8 | 6 | 2 | 3 | 7 | 5 | 4 | 7 | 7 | 7 |  |
|  |  | Ar | 6.54 | 3.79 | 3.59 | 5.41 | 4.54 | 1.99 | 2.45 | 4.4 | 3.48 | 3.73 | 5.22 | 5.22 | 5.21 |  |
|  |  | Ho | 0.77 | 0.77 | 0.33 | 0.82 | 0.76 | 0.2 | 0.27 | 0.62 | 0.52 | 0.63 | 0.92 | 0.72 | 0.72 |  |
|  |  | He | 0.75 | 0.65 | 0.43 | 0.8 | 0.67 | 0.36 | 0.38 | 0.65 | 0.52 | 0.61 | 0.79 | 0.71 | 0.73 |  |
|  |  | Fis | -0.04 | -0.2 | 0.26 | 0.01 | -0.13 | 0.35 | 0.3 | 0.08 | 0 | -0.08 | -0.17 | 0.02 | 0.04 |  |
|  |  | HWE | 0.83 | 0.33 | 1 | 0.89 | 0 | 0.42 | 1 | 0.01 | 0 | 0.7 | 0.06 | 0.83 | 0.91 |  |
|  | 11 - Scorff | N | 32 | 32 | 32 | 32 | 32 | 32 | 32 | 32 | 32 | 32 | 32 | 32 | 32 | 32 |
|  |  | A | 8 | 6 | 8 | 10 | 6 | 3 | 5 | 5 | 10 | 6 | 6 | 8 | 4 | 4 |
|  |  | Ar | 5.15 | 4.1 | 4.14 | 6.16 | 4.06 | 2.26 | 2.38 | 3.86 | 4.97 | 3.66 | 3.7 | 5.43 | 3.46 | 3.46 |
|  |  | Ho | 0.72 | 0.6 | 0.47 | 0.91 | 0.63 | 0.48 | 0.31 | 0.56 | 0.47 | 0.53 | 0.52 | 0.78 | 0.48 | 0.48 |
|  |  | He | 0.67 | 0.64 | 0.45 | 0.79 | 0.65 | 0.52 | 0.3 | 0.66 | 0.52 | 0.63 | 0.65 | 0.75 | 0.52 | 0.52 |
|  |  | Fis | -0.06 | 0.12 | 0.12 | -0.11 | 0.1 | 0.04 | 0.05 | 0.09 | 0.13 | 0.18 | 0.13 | -0.06 | 0.14 | 0.14 |
|  |  | HWE | 0.05 | 0.52 | 0.76 | 0.64 | 0.45 | 0.39 | 0.64 | 0.1 | 0.34 | 0.2 | 0.02 | 0.81 | 0.35 | 0.35 |
|  | 12 – Aulne | N | 14 | 14 | 14 | 14 | 14 | 14 | 14 | 14 | 14 | 14 | 14 | 14 | 14 | 14 |
|  |  | A | 6 | 5 | 4 | 7 | 4 | 2 | 2 | 3 | 8 | 3 | 5 | 5 | 4 | 4 |
|  |  | Ar | 4.8 | 3.96 | 2.71 | 5.69 | 3.93 | 2 | 1.94 | 2.76 | 6.23 | 2.97 | 4.67 | 4.59 | 3.14 | 3.14 |
|  |  | Ho | 0.62 | 0.57 | 0.21 | 0.83 | 0.5 | 0.54 | 0 | 0.29 | 0.42 | 0.67 | 0.44 | 0.77 | 0.57 | 0.57 |
|  |  | He | 0.69 | 0.52 | 0.21 | 0.76 | 0.73 | 0.51 | 0.17 | 0.32 | 0.69 | 0.63 | 0.67 | 0.76 | 0.52 | 0.52 |
|  |  | Fis | 0.11 | -0.11 | -0.04 | -0.1 | 0.33 | -0.06 | 1 | 0.11 | 0.41 | -0.07 | 0.35 | -0.01 | -0.1 | -0.1 |
|  |  | HWE | 0.26 | 0.85 | 1 | 0.27 | 0.06 | 0.79 | 0.05 | 0.35 | 0.05 | 0.69 | 0.11 | 0.5 | 0.75 | 0.75 |
|  | 13 – Trieux | N | 16 | 16 | 16 | 16 | 16 | 16 | 16 | 16 | 16 | 16 | 16 | 16 | 16 | 16 |
|  |  | A | 5 | 6 | 3 | 6 | 4 | 2 | 3 | 4 | 8 | 5 | 4 | 6 | 5 | 5 |
|  |  | Ar | 3.64 | 4.77 | 2.71 | 5.74 | 3.49 | 2 | 2.74 | 3.34 | 5.27 | 3.84 | 3.78 | 5.72 | 4.01 | 4.01 |
|  |  | Ho | 0.69 | 0.5 | 0.38 | 1 | 0.53 | 0.44 | 0.47 | 0.25 | 0.44 | 0.69 | 0.67 | 0.88 | 0.63 | 0.63 |
|  |  | He | 0.54 | 0.64 | 0.33 | 0.85 | 0.62 | 0.47 | 0.39 | 0.43 | 0.53 | 0.67 | 0.63 | 0.79 | 0.57 | 0.57 |
| Fis |  | -0.28 | 0.23 | -0.18 | -0.05 | 0.26 | 0.06 | -0.23 | 0.42 | 0.24 | 0.03 | -0.24 | -0.19 | 0.03 | 0.03 |  |
| HWE |  | 1 | 0.15 | 1 | 0.94 | 0.1 | 0.61 | 1 | 0.02 | 0.18 | 0.69 | 0.74 | 0.82 | 0.83 | 0.83 |  |
| Normandy | 15 – Vire | N | 29 | 29 | 29 | 29 | 29 | 29 | 29 | 29 | 29 | 29 | 29 | 29 | 29 |  |
|  |  | A | 6 | 4 | 4 | 7 | 6 | 2 | 2 | 6 | 5 | 5 | 6 | 8 | 5 |  |
|  |  | Ar | 4.84 | 3.35 | 3.13 | 4.6 | 3.83 | 1.99 | 1.81 | 4.1 | 2.86 | 4.17 | 4.96 | 5.86 | 3.62 |  |
|  |  | Ho | 0.83 | 0.52 | 0.45 | 0.69 | 0.43 | 0.31 | 0.17 | 0.55 | 0.14 | 0.66 | 0.76 | 0.83 | 0.69 |  |
|  |  | He | 0.77 | 0.6 | 0.38 | 0.65 | 0.46 | 0.35 | 0.16 | 0.63 | 0.26 | 0.66 | 0.76 | 0.79 | 0.56 |  |
|  |  | Fis | -0.08 | 0.14 | -0.17 | -0.07 | 0.17 | 0.13 | -0.08 | 0.12 | 0.47 | 0.01 | 0.01 | -0.05 | -0.23 |  |
|  |  | HWE | 0.83 | 0.33 | 1 | 0.89 | 0.06 | 0.42 | 1 | 0.01 | 0 | 0.7 | 0.06 | 0.82 | 0.91 |  |
|  | 16 – Orne | N | 8 | 8 | 8 | 8 | 8 | 8 | 8 | 8 | 8 | 8 | 8 | 8 | 8 | 8 |
|  |  | A | 4 | 5 | 4 | 5 | 5 | 2 | 3 | 4 | 4 | 4 | 5 | 7 | 4 | 4 |
|  |  | Ar | 4 | 4 | 4 | 4 | 5 | 2 | 3 | 4 | 4 | 3 | 5 | 6 | 4 | 4 |
|  |  | Ho | 0.38 | 0.38 | 0.63 | 0.5 | 0.75 | 0.38 | 0.38 | 0.75 | 0.38 | 0.63 | 1 | 0.75 | 0.38 | 0.38 |
|  |  | He | 0.68 | 0.35 | 0.65 | 0.73 | 0.67 | 0.33 | 0.43 | 0.74 | 0.59 | 0.49 | 0.8 | 0.62 | 0.69 | 0.69 |
|  |  | Fis | 0.33 | 0.07 | -0.06 | 0.23 | -0.26 | -0.08 | 0.13 | -0.01 | 0.54 | 0.16 | -0.07 | -0.09 | 0.48 | 0.48 |
|  |  | HWE | 0.08 | 1 | 0.57 | 0.06 | 0.91 | 1 | 0.38 | 0.44 | 0.03 | 1 | 1 | 1 | 0.09 | 0.09 |
| All | Overall | N | 301 | 301 | 301 | 301 | 301 | 301 | 301 | 301 | 301 | 301 | 301 | 301 | 301 |  |
|  |  | A | 21 | 10 | 15 | 14 | 14 | 4 | 7 | 9 | 19 | 7 | 10 | 11 | 9 |  |
|  |  | Ar | 6.51 | 4.46 | 4.16 | 6.86 | 4.11 | 2.18 | 2.28 | 4.09 | 5.57 | 4.26 | 5.59 | 5.68 | 4.91 |  |
|  |  | Ho | 0.71 | 0.58 | 0.46 | 0.76 | 0.6 | 0.39 | 0.34 | 0.54 | 0.5 | 0.63 | 0.72 | 0.76 | 0.68 |  |
|  |  | He | 0.78 | 0.68 | 0.45 | 0.81 | 0.65 | 0.46 | 0.37 | 0.65 | 0.64 | 0.64 | 0.77 | 0.75 | 0.72 |  |

**Supplementary Table S3:** Genetic diversity observed in *A. fallax* at each microsatellite markers (1 column = 1 markers). Details are displayed for each river (rows) and contains the following information: N = Number of individuals Sampled, A = Number of Alleles, Ar = Allelic Richness, Ho = Observed Heterozygosity, He = Expected Heterozygosity, Fis = Inbreeding coefficient, HWE = pvalue of being out of HWE.

| Region | River + ID |  | Alo01 | Alo06 | Alo07 | Alo09 | Alo15 | Alo16 | Alo17 | Alo26 | Alo29 | Alo32 | Alo33 | Alo43 | Alo45 |
| --- | --- | --- | --- | --- | --- | --- | --- | --- | --- | --- | --- | --- | --- | --- | --- |
| Spain | 1 -Minho | N | 24 | 24 | 24 | 24 | 24 | 24 | 24 | 24 | 24 | 24 | 24 | 24 | 24 |
|  |  | A | 4 | 4 | 5 | 5 | 6 | 1 | 6 | 6 | 4 | 5 | 3 | 4 | 4 |
|  |  | Ar | 2.8 | 3.09 | 3.73 | 4.5 | 4.4 | 1 | 4.52 | 4.99 | 3.93 | 4.58 | 2.81 | 3.23 | 2.51 |
|  |  | Ho | 0.55 | 0.42 | 0.46 | 0.71 | 0.55 | 0 | 0.83 | 0.75 | 0.65 | 0.58 | 0.46 | 0.38 | 0.21 |
|  |  | He | 0.46 | 0.41 | 0.5 | 0.74 | 0.59 | 0 | 0.75 | 0.78 | 0.74 | 0.75 | 0.48 | 0.42 | 0.2 |
|  |  | Fis | -0.21 | -0.04 | 0.13 | 0.04 | 0.08 | NA | -0.1 | 0.04 | 0.12 | 0.16 | 0.04 | 0.11 | -0.05 |
|  |  | HWE | 0.91 | 0.52 | 0.19 | 0.35 | 0.12 | No | 0.83 | 0.31 | 0.2 | 0.04 | 0.11 | 0.32 | 1 |
|  | 2 - Ulla | N | 31 | 31 | 31 | 31 | 31 | 31 | 31 | 31 | 31 | 31 | 31 | 31 | 31 |
|  |  | A | 7 | 3 | 6 | 8 | 4 | 2 | 4 | 5 | 5 | 4 | 4 | 3 | 3 |
|  |  | Ar | 3.97 | 1.71 | 3.51 | 4.48 | 2.97 | 1.92 | 2.75 | 4.07 | 4.19 | 2.72 | 3.14 | 1.52 | 2.45 |
|  |  | Ho | 0.71 | 0.1 | 0.45 | 0.58 | 0.65 | 0.26 | 0.26 | 0.63 | 0.7 | 0.5 | 0.54 | 0.06 | 0.29 |
|  |  | He | 0.62 | 0.09 | 0.43 | 0.54 | 0.58 | 0.23 | 0.29 | 0.66 | 0.68 | 0.4 | 0.51 | 0.06 | 0.44 |
|  |  | Fis | -0.15 | -0.02 | -0.05 | -0.07 | -0.11 | -0.13 | 0.11 | 0.05 | -0.03 | -0.25 | -0.07 | -0.01 | 0.34 |
|  |  | HWE | 0.85 | 1 | 0.07 | 0.56 | 0.81 | 1 | 0.03 | 0.07 | 0.04 | 1 | 0.18 | 1 | 0.05 |
| Southern France | 4- Adour | N | 15 | 15 | 15 | 15 | 15 | 15 | 15 | 15 | 15 | 15 | 15 | 15 | 15 |
|  |  | A | 6 | 6 | 7 | 6 | 6 | 3 | 3 | 6 | 3 | 6 | 4 | 3 | 2 |
|  |  | Ar | 4.77 | 5.22 | 5.99 | 4.39 | 4.82 | 2.5 | 2.96 | 5.09 | 3.53 | 4.82 | 3.66 | 2.48 | 1.53 |
|  |  | Ho | 0.73 | 0.53 | 0.67 | 0.6 | 0.4 | 0.07 | 0.4 | 0.43 | 0.6 | 0.73 | 0.55 | 0.08 | 0.07 |
|  |  | He | 0.71 | 0.75 | 0.8 | 0.57 | 0.69 | 0.3 | 0.63 | 0.74 | 0.71 | 0.69 | 0.64 | 0.22 | 0.07 |
|  |  | Fis | 0.01 | 0.24 | 0.11 | -0.08 | 0.48 | 0.78 | 0.45 | 0.36 | 0.18 | -0.04 | 0.15 | 0.66 | 0 |
|  |  | HWE | 0.71 | 0.02 | 0.11 | 0.48 | 0 | 0 | 0.04 | 0 | 0.25 | 0.65 | 0.35 | 0.04 | No |
| Atlantic | 6 - Dordogne | N | 15 | 15 | 15 | 15 | 15 | 15 | 15 | 15 | 15 | 15 | 15 | 15 | 15 |
|  |  | A | 5 | 5 | 6 | 6 | 4 | 2 | 4 | 4 | 7 | 5 | 5 | 4 | 1 |
|  |  | Ar | 4.48 | 4.32 | 4.77 | 5.16 | 3.32 | 1.99 | 2.93 | 3.48 | 2.53 | 3.44 | 3.89 | 2.86 | 1 |
|  |  | Ho | 0.73 | 0.33 | 0.67 | 0.64 | 0.38 | 0.21 | 0.77 | 0.47 | 0.67 | 0.27 | 0.64 | 0.13 | 0 |
|  |  | He | 0.69 | 0.46 | 0.72 | 0.78 | 0.54 | 0.39 | 0.7 | 0.59 | 0.72 | 0.68 | 0.56 | 0.19 | 0 |
|  |  | Fis | -0.06 | 0.28 | 0.07 | 0.18 | 0.3 | 0.46 | -0.1 | 0.22 | 0.07 | 0.62 | -0.15 | 0.32 | NA |
|  |  | HWE | 0.69 | 0.1 | 0.05 | 0.03 | 0.02 | 0.33 | 0.96 | 0.37 | 0.58 | 0.03 | 0.72 | 0.2 | No |
| Normandy | 16 – Orne | N | 15 | 15 | 15 | 15 | 15 | 15 | 15 | 15 | 15 | 15 | 15 | 15 | 15 |
|  |  | A | 4 | 4 | 2 | 5 | 8 | 2 | 4 | 6 | 4 | 3 | 4 | 4 | 3 |
|  |  | Ar | 3.46 | 3.23 | 1.79 | 3.82 | 5.83 | 1.96 | 3.51 | 4.98 | 3.07 | 2.79 | 3.57 | 3.9 | 2.07 |
|  |  | Ho | 0.6 | 0.27 | 0.13 | 0.4 | 0.87 | 0.13 | 0.6 | 0.67 | 0.47 | 0.53 | 0.62 | 0.53 | 0.13 |
|  |  | He (nb) | 0.49 | 0.36 | 0.13 | 0.45 | 0.71 | 0.24 | 0.59 | 0.78 | 0.54 | 0.56 | 0.6 | 0.68 | 0.13 |
|  |  | Fis | -0.24 | 0.26 | -0.04 | 0.12 | -0.22 | 0.45 | -0.02 | 0.15 | 0.15 | 0.05 | -0.02 | 0.22 | -0.02 |
|  |  | HWE | 1 | 0.13 | 1 | 0.23 | 0.99 | 0.2 | 0.56 | 0.26 | 0.34 | 0.53 | 0.63 | 0.04 | 1 |
| Mediterranean Sea – Rhone Watershed | 18 – Vidourle | N | 9 | 9 | 9 | 9 | 9 | 9 | 9 | 9 | 9 | 9 | 9 | 9 | 9 |
|  |  | A | 8 | 7 | 7 | 6 | 3 | 3 | 5 | 3 | 3 | 3 | 4 | 6 | 4 |
|  |  | Ar | 7.33 | 6.77 | 6.44 | 6 | 3 | 2.88 | 4.77 | 2.99 | 2.89 | 2.89 | 3.99 | 5.77 | 3.77 |
|  |  | Ho | 1 | 0.67 | 0.56 | 0.63 | 0.5 | 0.33 | 0.67 | 0.11 | 0.78 | 0.56 | 0.56 | 0.78 | 0.33 |
|  |  | He | 0.8 | 0.86 | 0.69 | 0.8 | 0.43 | 0.31 | 0.61 | 0.46 | 0.58 | 0.45 | 0.69 | 0.77 | 0.4 |
|  |  | Fis | -0.27 | 0.24 | 0.2 | 0.23 | -0.25 | -0.09 | -0.09 | 0.77 | -0.37 | -0.25 | 0.2 | -0.01 | 0.17 |
|  |  | HWE | 1 | 0.05 | 0.26 | 0.09 | 1 | 1 | 0.85 | 0.01 | 0.95 | 1 | 0.31 | 0.69 | 0.34 |
|  | 19 – Aude | N | 15 | 15 | 15 | 15 | 15 | 15 | 15 | 15 | 15 | 15 | 15 | 15 | 15 |
|  |  | A | 4 | 6 | 5 | 8 | 5 | 2 | 5 | 5 | 5 | 2 | 4 | 7 | 4 |
|  |  | Ar | 3.99 | 5.03 | 4.39 | 6.37 | 4.14 | 2 | 3.77 | 3.6 | 4.39 | 2 | 3.57 | 5.88 | 3.23 |
|  |  | Ho | 0.5 | 0.6 | 0.53 | 0.8 | 0.4 | 0.4 | 0.4 | 0.53 | 0.79 | 0.47 | 0.79 | 0.8 | 0.4 |
|  |  | He | 0.75 | 0.73 | 0.64 | 0.8 | 0.69 | 0.4 | 0.41 | 0.48 | 0.74 | 0.48 | 0.7 | 0.82 | 0.36 |
|  |  | Fis | 0.34 | 0.19 | 0.18 | 0 | 0.43 | 0.01 | 0.02 | -0.11 | -0.07 | 0.03 | -0.14 | 0.03 | -0.13 |
|  |  | HWE | 0.03 | 0.22 | 0.13 | 0.51 | 0 | 0.72 | 0.6 | 0.82 | 0.75 | 0.67 | 0.71 | 0.44 | 1 |
|  | 20 – Rhône1 | N | 30 | 30 | 30 | 30 | 30 | 30 | 30 | 30 | 30 | 30 | 30 | 30 | 30 |
|  |  | A | 9 | 9 | 7 | 12 | 8 | 3 | 6 | 7 | 6 | 5 | 6 | 9 | 4 |
|  |  | Ar | 5.04 | 6.02 | 4.39 | 8.46 | 4.31 | 2.07 | 3.86 | 4.37 | 3.91 | 3.49 | 4.36 | 6.47 | 2.53 |
|  |  | Ho | 0.73 | 0.73 | 0.5 | 0.7 | 0.8 | 0.2 | 0.47 | 0.37 | 0.44 | 0.73 | 0.76 | 0.97 | 0.27 |
|  |  | He | 0.75 | 0.72 | 0.5 | 0.89 | 0.69 | 0.19 | 0.45 | 0.54 | 0.57 | 0.61 | 0.72 | 0.84 | 0.24 |
|  |  | Fis | 0.03 | - | 0 | 0.21 | - | -0.08 | -0.04 | 0.32 | 0.23 | - | - | - | -0.09 |

|  |  | 0.02 |  |  |  | 0.17 |  |  |  | 0.21 |  |  |  | 0.06 | 0.15 |
| --- | --- | --- | --- | --- | --- | --- | --- | --- | --- | --- | --- | --- | --- | --- | --- |
|  |  | HWE | 0.48 | 0.1 | 0.64 | 0.02 | 0.88 | 1 | 0.74 | 0.02 | 0.03 | 0.96 | 0.8 | 0.83 | 1 |
|  | 21 – Rhône2 | N | 37 | 37 | 37 | 37 | 37 | 37 | 37 | 37 | 37 | 37 | 37 | 37 | 37 |
|  |  | A | 10 | 9 | 6 | 13 | 6 | 4 | 6 | 7 | 6 | 4 | 7 | 8 | 5 |
|  |  | Ar | 5.12 | 5.74 | 3.8 | 7.64 | 3.98 | 2.49 | 3.56 | 4.43 | 3.67 | 3.05 | 4.69 | 6.01 | 3.25 |
|  |  | Ho | 0.59 | 0.65 | 0.43 | 0.74 | 0.57 | 0.24 | 0.38 | 0.61 | 0.5 | 0.59 | 0.61 | 0.84 | 0.44 |
|  |  | He | 0.72 | 0.74 | 0.42 | 0.85 | 0.62 | 0.23 | 0.39 | 0.65 | 0.6 | 0.55 | 0.75 | 0.79 | 0.47 |
|  |  | Fis | 0.18 | 0.13 | -0.03 | 0.12 | 0.07 | -0.08 | 0.01 | 0.07 | 0.17 | -0.09 | 0.18 | -0.06 | 0.05 |
|  |  | HWE | 0.13 | **0.01** | 0.34 | **0** | **0.01** | 1 | 0.58 | 0.37 | 0.07 | 0.82 | **0** | 0.63 | 0.52 |
| Mediterranean Sea – Corsica | 22 - Tavignano1 | N | 50 | 50 | 50 | 50 | 50 | 50 | 50 | 50 | 50 | 50 | 50 | 50 | 50 |
|  |  | A | 5 | 6 | 6 | 7 | 7 | 2 | 4 | 5 | 2 | 3 | 6 | 3 | 4 |
|  |  | Ar | 3.33 | 3.87 | 5.03 | 4.89 | 3.72 | 1.97 | 3.17 | 4.39 | 1.16 | 2.98 | 3.84 | 2.82 | 3.56 |
|  |  | Ho | 0.54 | 0.51 | 0.82 | 0.51 | 0.45 | 0.31 | 0.65 | 0.78 | 0.02 | 0.58 | 0.64 | 0.44 | 0.66 |
|  |  | He | 0.55 | 0.55 | 0.77 | 0.71 | 0.52 | 0.29 | 0.66 | 0.74 | 0.02 | 0.64 | 0.65 | 0.43 | 0.68 |
|  |  | Fis | 0.01 | 0.07 | -0.06 | 0.28 | 0.14 | -0.06 | 0.02 | -0.05 | 0 | 0.09 | 0.02 | -0.02 | 0.02 |
|  |  | HWE | 0.53 | 0.01 | 0.85 | **0** | 0.21 | 0.81 | 0.37 | 0.8 | No | 0.31 | 0.53 | 0.71 | 0.37 |
|  | 23 Tavignano2 | N | 15 | 15 | 15 | 15 | 15 | 15 | 15 | 15 | 15 | 15 | 15 | 15 | 15 |
|  |  | A | 3 | 4 | 5 | 6 | 6 | 2 | 3 | 4 | 2 | 4 | 5 | 3 | 4 |
|  |  | Ar | 2.96 | 3.29 | 4.2 | 5.14 | 5.11 | 2 | 2.99 | 3.93 | 1.53 | 3.51 | 4.32 | 2.44 | 3.5 |
|  |  | Ho | 0.6 | 0.47 | 0.47 | 0.6 | 0.73 | 0.53 | 0.67 | 0.87 | 0.07 | 0.6 | 0.67 | 0.27 | 0.53 |
|  |  | He | 0.51 | 0.4 | 0.54 | 0.74 | 0.66 | 0.4 | 0.62 | 0.7 | 0.07 | 0.59 | 0.72 | 0.25 | 0.66 |
|  |  | Fis | -0.17 | -0.17 | 0.14 | 0.2 | -0.12 | -0.33 | -0.08 | -0.25 | 0 | -0.02 | 0.08 | -0.09 | 0.19 |
|  |  | HWE | 1 | 1 | 0.21 | 0.33 | 0.68 | 0.51 | 1 | 0.42 | No info | 0.35 | 0.38 | 1 | 0.05 |
| Overall | OverAll | N | 255 | 255 | 255 | 255 | 255 | 255 | 255 | 255 | 255 | 255 | 255 | 255 | 255 |
|  |  | A | 18 | 12 | 12 | 14 | 16 | 4 | 9 | 11 | 8 | 8 | 8 | 9 | 9 |
|  |  | Ar | 5.43 | 5.84 | 6.02 | 7.17 | 7.47 | 2.11 | 4.43 | 5.8 | 4.54 | 4.34 | 4.89 | 4.95 | 3.25 |
|  |  | Ho | 0.66 | 0.48 | 0.52 | 0.63 | 0.57 | 0.24 | 0.55 | 0.56 | 0.52 | 0.56 | 0.62 | 0.48 | 0.3 |
|  |  | He | 0.9 | 1 | 0.26 | 0.07 | 0.33 | 1 | 0.76 | 0.92 | No | 0.71 | 0.41 | 1 | 0.3 |

**Supplementary Table S4:** Linear model results for comparison among *A. alosa* population and among *A. fallax* population

|  |  | AIC | BIC | Log Likelihood | deviance | chisq | P |
| --- | --- | --- | --- | --- | --- | --- | --- |
| *A. alosa* | nul model | 465.33 | 474.72 | -229.66 | 459.33 |  |  |
| | Allelic Richness ~ River + (1 \|Loci) | 456.59 | 503.54 | -213.3 | 426.59 | 32.736 | 0.001064 |
| *A. fallax* | nul model | 461.32 | 470.21 | -227.66 | 455.32 |  |  |
| | Allelic Richness ~ River + (1 \|Loci) | 445.27 | 483.79 | -209.64 | 419.27 | 36.044 | 8.27E-05 |

**Supplementary Table S5:** Results of Tukey HSD test for differences in Allelic Richness among populations.

**A) *A. alosa***

| | Estimate | Std. Error | z value | Pr(>\|z\|) |
| --- | --- | --- | --- | --- |
| Aulne - Adour == 0 | -0.4523846 | 0.294764 | -1.535 | 0.9483 |
| Charente - Adour == 0 | 0.3633846 | 0.294764 | 1.233 | 0.9913 |
| Dordogne - Adour == 0 | 0.4812308 | 0.294764 | 1.633 | 0.9203 |
| Garonne - Adour == 0 | 0.3632308 | 0.294764 | 1.232 | 0.9913 |
| Loire - Adour == 0 | 0.4883846 | 0.294764 | 1.657 | 0.9115 |
| Minõ - Adour == 0 | 0.3676923 | 0.294764 | 1.247 | 0.9904 |
| Nivelle - Adour == 0 | -0.3055385 | 0.294764 | -1.037 | 0.9983 |
| Orne - Adour == 0 | -0.2512308 | 0.294764 | -0.852 | 0.9998 |
| Scorff - Adour == 0 | -0.1496154 | 0.294764 | -0.508 | 1 |
| Trioux - Adour == 0 | -0.3252308 | 0.294764 | -1.103 | 0.9968 |
| Vilaine - Adour == 0 | 0.0237692 | 0.294764 | 0.081 | 1 |
| Vire - Adour == 0 | -0.4738462 | 0.294764 | -1.608 | 0.9283 |
| Charente - Aulne == 0 | 0.8157692 | 0.294764 | 2.768 | 0.2178 |
| Dordogne - Aulne == 0 | 0.9336154 | 0.294764 | 3.167 | 0.0764 |
| Garonne - Aulne == 0 | 0.8156154 | 0.294764 | 2.767 | 0.2181 |
| Loire - Aulne == 0 | 0.9407692 | 0.294764 | 3.192 | 0.0723 |
| Minõ - Aulne == 0 | 0.8200769 | 0.294764 | 2.782 | 0.2106 |
| Nivelle - Aulne == 0 | 0.1468462 | 0.294764 | 0.498 | 1 |
| Orne - Aulne == 0 | 0.2011538 | 0.294764 | 0.682 | 1 |
| Scorff - Aulne == 0 | 0.3027692 | 0.294764 | 1.027 | 0.9984 |
| Trioux - Aulne == 0 | 0.1271538 | 0.294764 | 0.431 | 1 |
| Vilaine - Aulne == 0 | 0.4761538 | 0.294764 | 1.615 | 0.9254 |
| Vire - Aulne == 0 | -0.0214615 | 0.294764 | -0.073 | 1 |
| Dordogne - Charente == 0 | 0.1178462 | 0.294764 | 0.4 | 1 |
| Garonne - Charente == 0 | -0.0001538 | 0.294764 | -0.001 | 1 |
| Loire - Charente == 0 | 0.125 | 0.294764 | 0.424 | 1 |
| Minõ - Charente == 0 | 0.0043077 | 0.294764 | 0.015 | 1 |
| Nivelle - Charente == 0 | -0.6689231 | 0.294764 | -2.269 | 0.5399 |
| Orne - Charente == 0 | -0.6146154 | 0.294764 | -2.085 | 0.6756 |
| Scorff - Charente == 0 | -0.513 | 0.294764 | -1.74 | 0.8783 |
| Trioux - Charente == 0 | -0.6886154 | 0.294764 | -2.336 | 0.4901 |
| Vilaine - Charente == 0 | -0.3396154 | 0.294764 | -1.152 | 0.9952 |
| Vire - Charente == 0 | -0.8372308 | 0.294764 | -2.84 | 0.1827 |
| Garonne - Dordogne == 0 | -0.118 | 0.294764 | -0.4 | 1 |
| Loire - Dordogne == 0 | 0.0071538 | 0.294764 | 0.024 | 1 |
| Minõ - Dordogne == 0 | -0.1135385 | 0.294764 | -0.385 | 1 |
| Nivelle - Dordogne == 0 | -0.7867692 | 0.294764 | -2.669 | 0.2684 |
| Orne - Dordogne == 0 | -0.7324615 | 0.294764 | -2.485 | 0.384 |
| Scorff - Dordogne == 0 | -0.6308462 | 0.294764 | -2.14 | 0.6359 |
| Trieux - Dordogne == 0 | -0.8064615 | 0.294764 | -2.736 | 0.2329 |
| Vilaine - Dordogne == 0 | -0.4574615 | 0.294764 | -1.552 | 0.9442 |
| Vire - Dordogne == 0 | -0.9550769 | 0.294764 | -3.24 | 0.0629 |
| Loire - Garonne == 0 | 0.1251538 | 0.294764 | 0.425 | 1 |
| Minõ - Garonne == 0 | 0.0044615 | 0.294764 | 0.015 | 1 |
| Nivelle - Garonne == 0 | -0.6687692 | 0.294764 | -2.269 | 0.5416 |
| Orne - Garonne == 0 | -0.6144615 | 0.294764 | -2.085 | 0.6746 |
| Scorff - Garonne == 0 | -0.5128462 | 0.294764 | -1.74 | 0.879 |
| Trieux - Garonne == 0 | -0.6884615 | 0.294764 | -2.336 | 0.4911 |
| Vilaine - Garonne == 0 | -0.3394615 | 0.294764 | -1.152 | 0.9953 |
| Vire - Garonne == 0 | -0.8370769 | 0.294764 | -2.84 | 0.1833 |
| Minõ - Loire == 0 | -0.1206923 | 0.294764 | -0.409 | 1 |
| Nivelle - Loire == 0 | -0.7939231 | 0.294764 | -2.693 | 0.2557 |
| Orne - Loire == 0 | -0.7396154 | 0.294764 | -2.509 | 0.3686 |
| Scorff - Loire == 0 | -0.638 | 0.294764 | -2.164 | 0.6176 |
| Trieux - Loire == 0 | -0.8136154 | 0.294764 | -2.76 | 0.2208 |
| Vilaine - Loire == 0 | -0.4646154 | 0.294764 | -1.576 | 0.9374 |
| Vire - Loire == 0 | -0.9622308 | 0.294764 | -3.264 | 0.0585 |
| Nivelle - Minõ == 0 | -0.6732308 | 0.294764 | -2.284 | 0.5285 |
| Orne - Minõ == 0 | -0.6189231 | 0.294764 | -2.1 | 0.6642 |
| Scorff - Minõ == 0 | -0.5173077 | 0.294764 | -1.755 | 0.8712 |
| Trieux - Minõ == 0 | -0.6929231 | 0.294764 | -2.351 | 0.4802 |
| Vilaine - Minõ == 0 | -0.3439231 | 0.294764 | -1.167 | 0.9947 |
| Vire - Minõ == 0 | -0.8415385 | 0.294764 | -2.855 | 0.1771 |
| Orne - Nivelle == 0 | 0.0543077 | 0.294764 | 0.184 | 1 |
| Scorff - Nivelle == 0 | 0.1559231 | 0.294764 | 0.529 | 1 |
| Trieux - Nivelle == 0 | -0.0196923 | 0.294764 | -0.067 | 1 |
| Vilaine - Nivelle == 0 | 0.3293077 | 0.294764 | 1.117 | 0.9965 |
| Vire - Nivelle == 0 | -0.1683077 | 0.294764 | -0.571 | 1 |
| Scorff - Orne == 0 | 0.1016154 | 0.294764 | 0.345 | 1 |
| Trieux - Orne == 0 | -0.074 | 0.294764 | -0.251 | 1 |
| Vilaine - Orne == 0 | 0.275 | 0.294764 | 0.933 | 0.9994 |
| Vire - Orne == 0 | -0.2226154 | 0.294764 | -0.755 | 0.9999 |
| Trieux - Scorff == 0 | -0.1756154 | 0.294764 | -0.596 | 1 |
| Vilaine - Scorff == 0 | 0.1733846 | 0.294764 | 0.588 | 1 |
| Vire - Scorff == 0 | -0.3242308 | 0.294764 | -1.1 | 0.9969 |
| Vilaine - Trieux == 0 | 0.349 | 0.294764 | 1.184 | 0.994 |
| Vire - Trieux == 0 | -0.1486154 | 0.294764 | -0.504 | 1 |
| Vire - Vilaine == 0 | -0.4976154 | 0.294764 | -1.688 | 0.8998 |

**B) A. fallax:**

| | Estimate | Std.Error | zvalue | Pr(>\|z\|) |
| --- | --- | --- | --- | --- |
| Aude - Adour == 0 | 0.04646 | 0.37277 | 0.125 | 1 |
| Dordogne - Adour == 0 | -0.58346 | 0.37277 | -1.565 | 0.897 |
| Mino - Adour == 0 | -0.43577 | 0.37277 | -1.169 | 0.9858 |
| Orne - Adour == 0 | -0.59762 | 0.37277 | -1.603 | 0.8815 |
| Rhône1 - Adour == 0 | 0.43631 | 0.37277 | 1.17 | 0.9856 |
| Rhône2 - Adour == 0 | 0.57923 | 0.37277 | 1.554 | 0.9012 |
| Tavignano1 - Adour == 0 | -0.54008 | 0.37277 | -1.449 | 0.9359 |
| Tavignano2 - Adour == 0 | -0.52531 | 0.37277 | -1.409 | 0.9465 |
| Ulla - Adour == 0 | -0.94946 | 0.37277 | -2.547 | 0.2766 |
| Vidourle - Adour == 0 | 0.59615 | 0.37277 | 1.599 | 0.8831 |
| Dordogne - Aude == 0 | -0.62992 | 0.37277 | -1.69 | 0.8414 |
| Mino - Aude == 0 | -0.48223 | 0.37277 | -1.294 | 0.9702 |
| Orne - Aude == 0 | -0.64408 | 0.37277 | -1.728 | 0.8213 |
| Rhône1 - Aude == 0 | 0.38985 | 0.37277 | 1.046 | 0.994 |
| Rhône2 - Aude == 0 | 0.53277 | 0.37277 | 1.429 | 0.9414 |
| Tavignano1- Aude == 0 | -0.58654 | 0.37277 | -1.573 | 0.8935 |
| Tavignano2 - Aude == 0 | -0.57177 | 0.37277 | -1.534 | 0.9086 |
| Ulla - Aude == 0 | -0.99592 | 0.37277 | -2.672 | 0.2132 |
| Vidourle - Aude == 0 | 0.54969 | 0.37277 | 1.475 | 0.9281 |
| Mino - Dordogne == 0 | 0.14769 | 0.37277 | 0.396 | 1 |
| Orne - Dordogne == 0 | -0.01415 | 0.37277 | -0.038 | 1 |
| Rhône1 - Dordogne == 0 | 1.01977 | 0.37277 | 2.736 | 0.1841 |
| Rhône2 - Dordogne == 0 | 1.16269 | 0.37277 | 3.119 | 0.0669 |
| Tavignano1- Dordogne == 0 | 0.04338 | 0.37277 | 0.116 | 1 |
| Tavignano2 - Dordogne == 0 | 0.05815 | 0.37277 | 0.156 | 1 |
| Ulla - Dordogne == 0 | -0.366 | 0.37277 | -0.982 | 0.9964 |
| Vidourle - Dordogne == 0 | 1.17962 | 0.37277 | 3.164 | 0.0584 |
| Orne - Mino == 0 | -0.16185 | 0.37277 | -0.434 | 1 |
| Rhône1 - Mino == 0 | 0.87208 | 0.37277 | 2.339 | 0.4078 |
| Rhône2 - Mino == 0 | 1.015 | 0.37277 | 2.723 | 0.1901 |
| Tavignano1- Mino == 0 | -0.10431 | 0.37277 | -0.28 | 1 |
| Tavignano2 - Mino == 0 | -0.08954 | 0.37277 | -0.24 | 1 |
| Ulla - Mino == 0 | -0.51369 | 0.37277 | -1.378 | 0.9539 |
| Vidourle - Mino == 0 | 1.03192 | 0.37277 | 2.768 | 0.1696 |
| Rhône1 - Orne == 0 | 1.03392 | 0.37277 | 2.774 | 0.168 |
| Rhône2 - Orne == 0 | 1.17685 | 0.37277 | 3.157 | 0.0604 |
| Tavignano1- Orne == 0 | 0.05754 | 0.37277 | 0.154 | 1 |
| Tavignano2 - Orne == 0 | 0.07231 | 0.37277 | 0.194 | 1 |
| Ulla - Orne == 0 | -0.35185 | 0.37277 | -0.944 | 0.9974 |
| Vidourle - Orne == 0 | 1.19377 | 0.37277 | 3.202 | 0.0528 |
| Rhône2 - Rhône1 == 0 | 0.14292 | 0.37277 | 0.383 | 1 |
| Tavignano1- Rhône1 == 0 | -0.97638 | 0.37277 | -2.619 | 0.2389 |
| Tavignano2 - Rhône1 == 0 | -0.96162 | 0.37277 | -2.58 | 0.2593 |
| Ulla - Rhône1 == 0 | -1.38577 | 0.37277 | -3.717 | <0.01 |
| Vidourle - Rhône1 == 0 | 0.15985 | 0.37277 | 0.429 | 1 |
| Tavignano1- Rhône2 == 0 | -1.11931 | 0.37277 | -3.003 | 0.0937 |
| Tavignano2 - Rhône2 == 0 | -1.10454 | 0.37277 | -2.963 | 0.1033 |
| Ulla - Rhône2 == 0 | -1.52869 | 0.37277 | -4.101 | <0.01 |
| Vidourle - Rhône2 == 0 | 0.01692 | 0.37277 | 0.045 | 1 |
| Tavignano2 - Tavignano1== 0 | 0.01477 | 0.37277 | 0.04 | 1 |
| Ulla - Tavignano1== 0 | -0.40938 | 0.37277 | -1.098 | 0.9912 |
| Vidourle - Tavignano1== 0 | 1.13623 | 0.37277 | 3.048 | 0.0822 |
| Ulla - Tavignano2 == 0 | -0.42415 | 0.37277 | -1.138 | 0.9884 |
| Vidourle - Tavignano2 == 0 | 1.12146 | 0.37277 | 3.008 | 0.0911 |
| Vidourle - Ulla == 0 | 1.54562 | 0.37277 | 4.146 | <0.01 |

**Supplementary Table S6: Mean Efficiency and Accuracy of hybrid identification for Structure and NewHybrid when considering different q-value threshold obtained on simulated data from HybridLab**

**Tq = Threshold q. value.**

**Efficiency:** Proportion of individuals in a group correctly identified. E.g. hybrid identification efficiency = number of hybrid individuals correctly identified as hybrids/total number of hybrids actually in the sample.

**Accuracy:** Proportion of an identified group that truly belongs to that category. E.g. hybrid identification accuracy = number of hybrid individuals in the hybrid group/total number of individuals in the group.

|  |  |  |  |  | Efficiency |  |  | Accuracy |  |  |
| --- | --- | --- | --- | --- | --- | --- | --- | --- | --- | --- |
| Tq | Method | N. simulate<br>d inds. | True Hybrid<br>Proportion<br>(%) | Observed<br>Hybrid<br>Proportion<br>(%) | Pure *A.*<br>*alosa* | Pure *A.*<br>*fallax* | Hybrids | Pure *A.*<br>*alosa* | Pure *A.*<br>*fallax* | Hybrid<br>s |
| 0.9 | Structure | 700 | 0 | 1.386 | 0.984 | 0.989 | - | 1 | 1 | - |
| 0.8 |  |  |  | 0.29 | 0.995 | 0.998 | - | 1 | 1 | - |
| 0.75 |  |  |  | 0.192 | 0.996 | 0.999 | - | 1 | 1 | - |
| 0.9 | New<br>Hybrids |  |  | 0.079 | 0.979 | 0.993 | - | 1 | 1 | - |
| 0.8 |  |  |  | 0.143 | 0.987 | 0.997 | - | 1 | 1 | - |
| 0.75 |  |  |  | 0.143 | 0.989 | 0.999 | - | 1 | 1 | - |
| 0.9 | Structure | 740 | 5 | 7.88 | 0.968 | 0.980 | 0.930 | 0.991 | 0.996 | 0.689 |
| 0.8 |  |  |  | 6.05 | 0.990 | 0.997 | 0.885 | 0.988 | 0.992 | 0.896 |
| 0.75 |  |  |  | 5.75 | 0.995 | 0.998 | 0.830 | 0.983 | 0.988 | 0.941 |
| 0.9 | New<br>Hybrids |  |  | 4.70 | 0.957 | 0.979 | 0.806 | 0.994 | 0.996 | 0.912 |
| 0.8 |  |  |  | 4.97 | 0.966 | 0.982 | 0.829 | 0.993 | 0.996 | 0.888 |
| 0.75 |  |  |  | 5.14 | 0.969 | 0.985 | 0.842 | 0.992 | 0.996 | 0.874 |

**Supplementary Table S7:** q-values of individuals classified as hybrids according to *New Hybrids* as well as q-value from Structure along with their 90% confidence intervals (available only in Structure).

Three categories were considered in *New hybrids* corresponding either to i) pure *A. alosa*, ii) pure *A. fallax* or iii) *hybrid*. Hybrids from new *NewHybrids* could correspond to either F1, F2 or backcrosses and were lumped into a single category given that the highest power and efficiency were reached when these categories were merged.

In Structure two categories were possible, corresponding to either *A. fallax* or *A. alosa*, with hybrid expect to displayed intermediate q-values (from 0.25 to 0.75 depending on the directions of introgression).

| Region | Population | Individual Id | New Hybrids q-values |  |  | Structure q-values |  |  |  |
| --- | --- | --- | --- | --- | --- | --- | --- | --- | --- |
|  |  |  | qA_ alosa | qA_ fallax | q_ hybrid | qA alosa | 90% CI | qA fallax | 90% CI |
| Atlantic (south) | Adour | ALFA40 | 0 | 0.523 | 0.477 | 0.112 | (0-0.285) | 0.888 | (0.715-1) |
| Spain | Ulla | AUPH06 | 0 | 0.023 | 0.977 | 0.358 | (0.156-0.574) | 0.642 | (0.426-0.844) |
| Atlantic | Pertuis Charentais | AAPC01 | 0 | 0 | 1 | 0.498 | (0.293-0.707) | 0.502 | (0.293-0.707) |
| Atlantic | Vienne | AAVi7 | 0 | 0.001 | 0.999 | 0.459 | (0.214-0.706) | 0.541 | (0.294-0.786) |
| Atlantic | Charente | AACH15 | 0 | 0 | 1 | 0.483 | (0.237-0.737) | 0.517 | (0.263-0.763) |
| Atlantic | Charente | AACH12 | 0.594 | 0 | 0.406 | 0.847 | (0.635-1) | 0.153 | (0-0.365) |
| Atlantic | Charente | AACH2 | 0.558 | 0 | 0.442 | 0.759 | (0.486-1) | 0.241 | (0-0.514) |
| Atlantic | Charente | AACH8 | 0.597 | 0 | 0.403 | 0.861 | (0.617-1) | 0.139 | (0-0.383) |
| Atlantic | Charente | AFCH298 | 0.056 | 0 | 0.944 | 0.697 | (0.482-0.889) | 0.303 | (0.111-0.518) |
| Atlantic | Loire Estuary | AAL1016 | 0 | 0 | 1 | 0.416 | (0.205-0.638) | 0.584 | (0.362-0.795) |
| Atlantic | Loire Estuary | AAL1020 | 0.419 | 0 | 0.581 | 0.836 | (0.643-1) | 0.164 | (0-0.357) |
| Atlantic | Loire Estuary | AAL1023 | 0.277 | 0 | 0.723 | 0.821 | (0.637-0.967) | 0.179 | (0.033-0.363) |
| Atlantic | Loire Estuary | AAL1024 | 0.341 | 0 | 0.659 | 0.786 | (0.554-1) | 0.214 | (0-0.446) |
| Atlantic | Loire | AAL52 | 0.258 | 0 | 0.742 | 0.823 | (0.623-0.98) | 0.177 | (0.02-0.377) |
| Atlantic | Loire | AAL54 | 0.001 | 0 | 1 | 0.623 | (0.406-0.822) | 0.377 | (0.178-0.594) |
| Atlantic | Loire | AAL167 | 0.618 | 0 | 0.382 | 0.865 | (0.609-1) | 0.135 | (0-0.391) |
| Brittany | Scorff | AASC2431 | 0.483 | 0 | 0.517 | 0.826 | (0.597-1) | 0.174 | (0-0.403) |
| Brittany | Scorff | AASC2445 | 0.053 | 0 | 0.947 | 0.74 | (0.538-0.914) | 0.26 | (0.086-0.462) |
| Brittany | South Brittany | AA0154 | 0 | 0 | 1 | 0.5 | (0.263-0.731) | 0.5 | (0.269-0.737) |
| Brittany | South Brittany | AA0155 | 0 | 0 | 1 | 0.501 | (0.277-0.722) | 0.499 | (0.278-0.723) |
| Brittany | South Brittany | AA161 | 0.232 | 0 | 0.768 | 0.672 | (0.406-0.962) | 0.328 | (0.038-0.594) |
| Brittany | South Brittany | AFSB94 | 0.009 | 0 | 0.991 | 0.616 | (0.391-0.825) | 0.384 | (0.175-0.609) |
| Normandy | Tamar | AATAM2 | 0.005 | 0 | 0.995 | 0.441 | (0.215-0.68) | 0.559 | (0.32-0.785) |
| Normandy | Calais | ALFA99 | 0 | 0.314 | 0.686 | 0.246 | (0.016-0.488) | 0.754 | (0.512-0.984) |

**Supplementary Table S8:** Pairwise weir & cockerham $F_{ST}$ between all populations and between-species. Sorted by species and from South to North. Between-species comparisons are in grey.

|  | Alosa alosa |  |  |  |  |  |  |  |  |  |  |  |  | A. fallax fallax |  |  |  |  | A. agone/A fallax rhodanensis |  |  |  | A. fallax sp. |
| --- | --- | --- | --- | --- | --- | --- | --- | --- | --- | --- | --- | --- | --- | --- | --- | --- | --- | --- | --- | --- | --- | --- | --- |
|  | Minho | Nivelle | Adour | Dordogne | Garonne | Charente | Loire | Vilaine | Scorff | Aulne | Trieux | Vire | Orne | Minho | Ulla | Adour | Dordogne | Orne | Vidourle | Aude | Rhône1 | Rhône2 | Tavignano1 |
| Nivelle | 0.137 | 0 |  |  |  |  |  |  |  |  |  |  |  |  |  |  |  |  |  |  |  |  |  |
| Adour | 0.118 | 0.065 | 0 |  |  |  |  |  |  |  |  |  |  |  |  |  |  |  |  |  |  |  |  |
| Dordogne | 0.075 | 0.061 | 0.015 | 0 |  |  |  |  |  |  |  |  |  |  |  |  |  |  |  |  |  |  |  |
| Garonne | 0.074 | 0.07 | 0.013 | -0.001 | 0 |  |  |  |  |  |  |  |  |  |  |  |  |  |  |  |  |  |  |
| Charente | 0.085 | 0.095 | 0.03 | 0.003 | 0.006 | 0 |  |  |  |  |  |  |  |  |  |  |  |  |  |  |  |  |  |
| Loire | 0.064 | 0.06 | 0.031 | 0.005 | 0.011 | 0.018 | 0 |  |  |  |  |  |  |  |  |  |  |  |  |  |  |  |  |
| Vilaine | 0.092 | 0.059 | 0.027 | 0.018 | 0.024 | 0.031 | 0.005 | 0 |  |  |  |  |  |  |  |  |  |  |  |  |  |  |  |
| Scorf | 0.085 | 0.086 | 0.041 | 0.03 | 0.043 | 0.056 | 0.027 | 0.024 | 0 |  |  |  |  |  |  |  |  |  |  |  |  |  |  |
| Aulne | 0.122 | 0.117 | 0.056 | 0.059 | 0.053 | 0.05 | 0.045 | 0.05 | 0.031 | 0 |  |  |  |  |  |  |  |  |  |  |  |  |  |
| Trieux | 0.108 | 0.092 | 0.039 | 0.045 | 0.046 | 0.046 | 0.029 | 0.022 | 0.007 | 0.008 | 0 |  |  |  |  |  |  |  |  |  |  |  |  |
| Vire | 0.14 | 0.08 | 0.051 | 0.062 | 0.049 | 0.072 | 0.044 | 0.042 | 0.05 | 0.06 | 0.046 | 0 |  |  |  |  |  |  |  |  |  |  |  |
| Orne | 0.11 | 0.071 | 0.038 | 0.039 | 0.028 | 0.048 | 0.04 | 0.038 | 0.038 | 0.049 | 0.044 | 0.031 | 0 |  |  |  |  |  |  |  |  |  |  |
| Minho | 0.361 | 0.333 | 0.33 | 0.302 | 0.321 | 0.3 | 0.319 | 0.313 | 0.331 | 0.358 | 0.349 | 0.351 | 0.335 | 0 |  |  |  |  |  |  |  |  |  |
| Ulla | 0.442 | 0.401 | 0.42 | 0.383 | 0.401 | 0.376 | 0.394 | 0.395 | 0.409 | 0.449 | 0.437 | 0.431 | 0.423 | 0.147 | 0 |  |  |  |  |  |  |  |  |
| Adour | 0.314 | 0.301 | 0.287 | 0.263 | 0.281 | 0.26 | 0.275 | 0.274 | 0.292 | 0.297 | 0.301 | 0.316 | 0.291 | 0.081 | 0.204 | 0 |  |  |  |  |  |  |  |
| Dordogne | 0.379 | 0.354 | 0.352 | 0.32 | 0.34 | 0.313 | 0.332 | 0.333 | 0.349 | 0.357 | 0.36 | 0.375 | 0.363 | 0.088 | 0.233 | 0.029 | 0 |  |  |  |  |  |  |
| Orne | 0.394 | 0.356 | 0.363 | 0.33 | 0.35 | 0.33 | 0.343 | 0.349 | 0.36 | 0.384 | 0.374 | 0.385 | 0.375 | 0.207 | 0.289 | 0.18 | 0.144 | 0 |  |  |  |  |  |
| Vidouroule | 0.308 | 0.283 | 0.286 | 0.256 | 0.281 | 0.274 | 0.278 | 0.287 | 0.297 | 0.336 | 0.325 | 0.34 | 0.287 | 0.245 | 0.29 | 0.177 | 0.24 | 0.217 | 0 |  |  |  |  |
| Aude | 0.298 | 0.279 | 0.273 | 0.245 | 0.272 | 0.261 | 0.267 | 0.275 | 0.284 | 0.314 | 0.305 | 0.321 | 0.273 | 0.224 | 0.253 | 0.169 | 0.217 | 0.17 | 0.011 | 0 |  |  |  |
| Rhône1 | 0.316 | 0.292 | 0.287 | 0.265 | 0.286 | 0.275 | 0.282 | 0.286 | 0.301 | 0.326 | 0.316 | 0.327 | 0.286 | 0.229 | 0.253 | 0.182 | 0.22 | 0.173 | 0.001 | 0.01 | 0 |  |  |
| Rhône2 | 0.318 | 0.282 | 0.278 | 0.259 | 0.28 | 0.274 | 0.278 | 0.281 | 0.296 | 0.323 | 0.312 | 0.318 | 0.282 | 0.227 | 0.264 | 0.184 | 0.225 | 0.166 | 0.016 | 0.011 | -0.002 | 0 |  |
| Tavignano1 | 0.361 | 0.328 | 0.333 | 0.316 | 0.326 | 0.312 | 0.324 | 0.329 | 0.346 | 0.355 | 0.348 | 0.361 | 0.346 | 0.286 | 0.328 | 0.253 | 0.288 | 0.323 | 0.242 | 0.233 | 0.237 | 0.252 | 0 |
| Tavignano2 | 0.363 | 0.329 | 0.332 | 0.31 | 0.322 | 0.301 | 0.314 | 0.325 | 0.345 | 0.358 | 0.348 | 0.366 | 0.356 | 0.303 | 0.345 | 0.26 | 0.308 | 0.358 | 0.262 | 0.248 | 0.249 | 0.268 | 0.011 |

[illegible]

**Supplementary Table S9:** Mean membership assignment of *A. alosa* individuals averaged by river to each cluster found in the DAPC displayed in Fig 2c.

| Region | River | cluster1 | cluster2 | cluster3 | cluster4 | cluster5 | cluster6 |
| --- | --- | --- | --- | --- | --- | --- | --- |
| Spain | Minho | 0 | 0.1 | 0 | 0 | 0.05 | 0.85 |
| Southern France | Nivelle | 0 | 0.04 | 0.62 | 0.04 | 0.26 | 0.03 |
|  | Adour | 0.32 | 0.15 | 0.2 | 0.24 | 0.09 | 0 |
| Atlantic | Charente | 0.53 | 0.12 | 0.12 | 0.03 | 0.08 | 0.12 |
|  | Dordogne | 0.38 | 0.04 | 0.21 | 0.08 | 0.18 | 0.12 |
|  | Garonne | 0.3 | 0.19 | 0.14 | 0.06 | 0.21 | 0.1 |
|  | Vienne | 0.2 | 0.37 | 0.2 | 0 | 0.03 | 0.2 |
|  | Loire | 0.12 | 0.09 | 0.09 | 0.15 | 0.41 | 0.15 |
|  | Vilaine | 0.22 | 0.23 | 0.19 | 0.14 | 0.16 | 0.07 |
| Brittany | Scorff | 0.12 | 0.08 | 0.06 | 0.45 | 0.16 | 0.13 |
|  | Aulne | 0.22 | 0.15 | 0.07 | 0.48 | 0 | 0.07 |
|  | Trieux | 0.18 | 0.13 | 0.07 | 0.49 | 0.07 | 0.06 |
|  | Sélune | 0.19 | 0.4 | 0.21 | 0.01 | 0.19 | 0 |
| Normandy | Orne | 0 | 0.48 | 0.25 | 0.25 | 0 | 0.01 |
|  | Vire | 0.01 | 0.62 | 0.02 | 0.18 | 0.16 | 0 |

**Supplementary Table S10:** Mean membership assignment of *A. fallax* individuals averaged by river to each cluster found in the DAPC displayed in Fig 2d.

[illegible]

| **Region** | **RiverName** | **cluster**<br>**1** | **cluster**<br>**2** | **cluster**<br>**3** | **cluster**<br>**4** | **cluster**<br>**5** | **cluster**<br>**6** |
| --- | --- | --- | --- | --- | --- | --- | --- |
| Mediterranean<br>Sea – ‘Corsica’ | Tavignano_1 | **0.92** | 0 | 0 | 0.08 | 0 | 0 |
|  | Tavignano_2 | **1** | 0 | 0 | 0 | 0 | 0 |
| Mediterranean<br>Sea – Rhone | Rhône_1 | 0.03 | 0 | 0 | **0.97** | 0 | 0 |
|  | Rhône_2 | 0 | 0 | 0 | **1** | 0 | 0 |
|  | Vidourle | 0 | 0 | 0 | **1** | 0 | 0 |
|  | Aude | 0 | 0 | 0.27 | **0.73** | 0 | 0 |
| Spain | Ulla | 0 | 0 | 0 | 0 | 0.03 | **0.97** |
|  | Minho | 0 | 0 | 0 | 0 | **0.84** | 0.16 |
|  | Adour | 0 | 0.12 | 0.01 | 0 | **0.87** | 0 |
| Atlantic | Dordogne | 0 | **0.31** | 0.13 | 0 | **0.56** | 0 |
|  | Oleron | 0 | **0.5** | 0 | 0 | **0.5** | 0 |
|  | Pertuis_charentais | 0 | **0.63** | 0.26 | 0 | 0.11 | 0 |
|  | Charente | 0 | 0.25 | 0.25 | 0 | 0.25 | 0.25 |
|  | Loire | 0 | 0 | 0.14 | 0 | 0.86 | 0 |
|  | Loire Estuary | 0 | 0 | 0 | 0 | **1** | 0 |
|  | Vilaine | 0 | 0 | 0 | 0 | **1** | 0 |
| Brittany | South Brittany | 0 | 0 | 0 | 0 | **1** | 0 |
|  | South_Brittany | 0 | 0 | **0.82** | 0 | 0.18 | 0 |
| Normandy | Orne | 0 | 0.07 | **0.8** | 0 | 0.13 | 0 |
|  | North Sea | 0 | 0.16 | **0.83** | 0 | 0 | 0 |
| Bay of Biscay | Bay_of_Biscay | 0.8 | 0 | 0.2 | 0 | 0 | 0 |

**Supplementary Table S11: Model choice for each species and lineages.** Posterior probabilities (P) of each model and its alternative were obtained through the neural network approach implemented in the ABC package. Values are display for the between-species comparison as well as between lineages of *A. fallax*. At the first round of comparison model with gene flow were compared against the model of strict isolation. At the second round, then the two model with ongoing gene-flow (IM and SC) were compared against model with Ancient gene-flow (AM). Finally, the two best model (IM and SC) were compared against each other at the last round. AM = Ancient Migration, IM = Isolation with Migration, SC = Secondary Contact SI = Strict Isolation.

|  |  | pairwise comparison |  |
| --- | --- | --- | --- |
|  |  | *A. Alosa* vs *A. fallax* “Atlantic” |  |
| posterior probabilities | Round1 | P(AM) = 0.72 | P(SI) = 0.28 |
|  |  | P(IM) = 0.851 | P(SI) = 0.149 |
|  |  | P(SC) = 0.980 | P(SI) = 0.02 |
|  | Round2 | P(IM) = 0.599 | P(AM) = 0.401 |
|  |  | P(SC) = 0.669 | P(AM) = 0.331 |
|  | Round3 | **P(SC) = 0.738** | P(IM) = 0.262 |
|  | *A. fallax* Mediterranean sea vs *A. fallax* Corsica |  |  |
|  | Round1 | P(AM) = 0.188 | P(SI) = 0.812 |
|  |  | P(IM) = 0.960 | P(SI) = 0.040 |
|  |  | P(SC) = 0.934 | P(SI) = 0.066 |
|  | Round2 | P(IM) = 0.948 | P(AM) = 0.052 |
|  |  | P(SC) = 0.935 | P(AM) = 0.065 |
|  | Round3 | **P(SC) = 0.517** | P(IM) = 0.482 |
|  | *A. fallax* Mediterranean sea vs *A. fallax* Corsica |  |  |
|  | Round1 | P(AM) = 0.415 | P(SI) = 0.585 |
|  |  | P(IM) = 0.981 | P(SI) = 0.02 |
|  |  | P(SC) = 0.964 | P(SI) = 0.036 |
|  | Round2 | P(SC) = 0.801 | P(AM) = 0.199 |
|  |  | P(IM) = 0.877 | P(AM) = 0.123 |
|  | Round3 | **P(SC) = 0.779** | P(IM) = 0.221 |

**Supplementary Table S12:** Prior and posterior parameter estimates for ABC computations. Uniform prior (U) distribution were used.

The values displayed for the rate of migration are scale by $4 \cdot N_{\text{ref}} \cdot \mu$ assuming $N_{\text{ref}} = 50,000$ and $\mu = 2.5 \times 10^{-4}$ mutations/bp/generation, as coalescent simulation are always scaled in the coalescent simulator used (ms).

$N_{\text{allis\_shad}}$ = effective population size of Allis shad (*A. alosa*).

$N_{\text{twaitte\_shad}}$ = effective population size of twaitte shad (*A. fallax*)

$m_1$ = effective migration rate from population 2 into population 1,

$m_2$ = effective migration rate population 1 into population 2. Here population 1 = *A. alosa* and population 2 = *A. fallax*.

$T_{\text{split}}$ = Divergence Time between species (in generations) and $T_{\text{sc}}$ = Time of secondary contact between the two species.

|  |  | Ne *A. alosa* | Ne *A. fallax* | Ne ancestral | m1 | m2 | Tsc | Tsplit |
| --- | --- | --- | --- | --- | --- | --- | --- | --- |
| Comparis on | prior | U[0-500,000] | U[0-500,000] | U[0-2,500,000] | U[0-15] | U[0-15] | U[0-Ts] | U[0-2,000,000] |
| between species | **2.5%CI** | 47 | 940 | 39,074 | 5.22E-06 | 2.26E-06 | 2,4432 | 119,909 |
|  | **median** | 209 | 2,573 | 617,465 | 4.48E-05 | 2.62E-05 | 15,6815 | 774,259 |
|  | **mean** | 314 | 3,097 | 629,502 | 4.24E-05 | 2.93E-05 | 21,1789 | 801,132 |
|  | **97.5%CI** | 646 | 5,649 | 1,228,609 | 7.25E-05 | 6.90E-05 | 62,1683 | 1,597,823 |
|  |  | Ne *A. fallax* "Atlantic" | Ne *A. fallax* "Mediterranean" | Nancestral | m1 | m2 | Tsc | Tsplit |
| A. fallax Atlantic vs A. fallax "mediterranean sea" | **2.5%CI** | 155 | 675 | 19485 | 5.54E-06 | 3.257E-06 | 15860 | 69,440 |
|  | **median** | 620 | 3,375 | 608270 | 4.13E-05 | 4.605E-05 | 163900 | 630,980 |
|  | **mean** | 1,640 | 4,510 | 610515 | 4.11E-05 | 4.360E-05 | 224100 | 710,740 |
|  | **97.5%CI** | 5,560 | 9,695 | 1211000 | 7.30E-05 | 7.376E-05 | 727800 | 970,320 |
|  |  | Ne *A. fallax* "mediterranean" | Ne *A. fallax* "Corsica" | Nancestral | m1 | m2 | Tsc | Tsplit |
| A. fallax "mediterranean sea" vs A. fallax Corsica | **2.5%CI** | 99 | 6,624 | 27,820 | 0.0000005 | 0.0000015 | 6317 | 34,496 |
|  | **median** | 348 | 19,508 | 620,842 | 0.0000092 | 0.0000274 | 57423 | 351,117 |
|  | **mean** | 392 | 22,467 | 619,242 | 0.0000146 | 0.0000303 | 99491 | 400,764 |
|  | **97.5%CI** | 861 | 51,808 | 120,6417 | 0.0000623 | 0.0000694 | 439084 | 959,376 |

**Fig S1: Extracted from Rougemont & Bernatchez 2018 (Evolution):** Representation of the demographic scenarios compared in this study: Strict Isolation (SI), Isolation with constant Migration (IM) Ancient Migration (AM) and Secondary Contact (SC). All models share the following parameters: $T_{\text{SPLIT}}$: number of generation of divergence (backwards in time). $N_{\text{anc}}$, $N_1$, $N_2$: effective population size of the ancestral population, of the first and second daughter population compared. $M_1$ and $M_2$ represent the effective migration rates per generation with $m$ the proportion of population made of migrants from the other populations. $T_{\text{AM}}$ is the number of generations since the two populations have diverged without gene flow. $T_{\text{SC}}$ is the number of generations since the populations have started exchanging alleles (secondary contact) after a period of isolation.

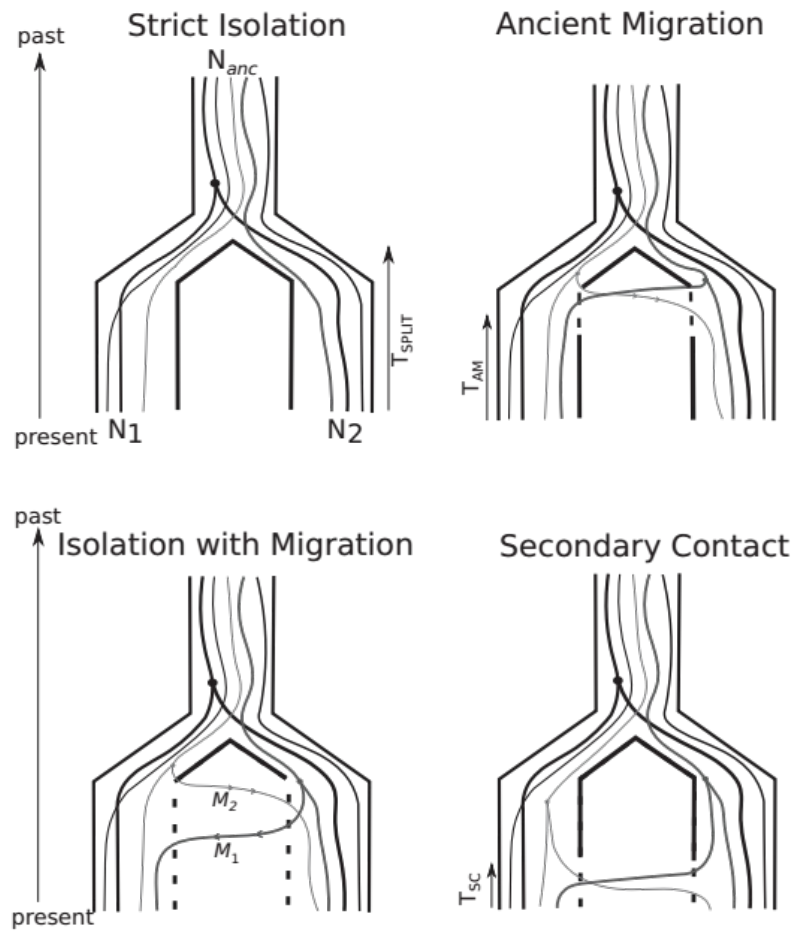

**Figure S2 : Results of Structure model choice.** A) Delta K for *A. alosa* obtained using Evanno et al. (2005) B) L(K) for *A. alosa* obtained using pritchard et al. (1999) method, C) Delta K for *A. fallax* and D) L(K) for *A. fallax*

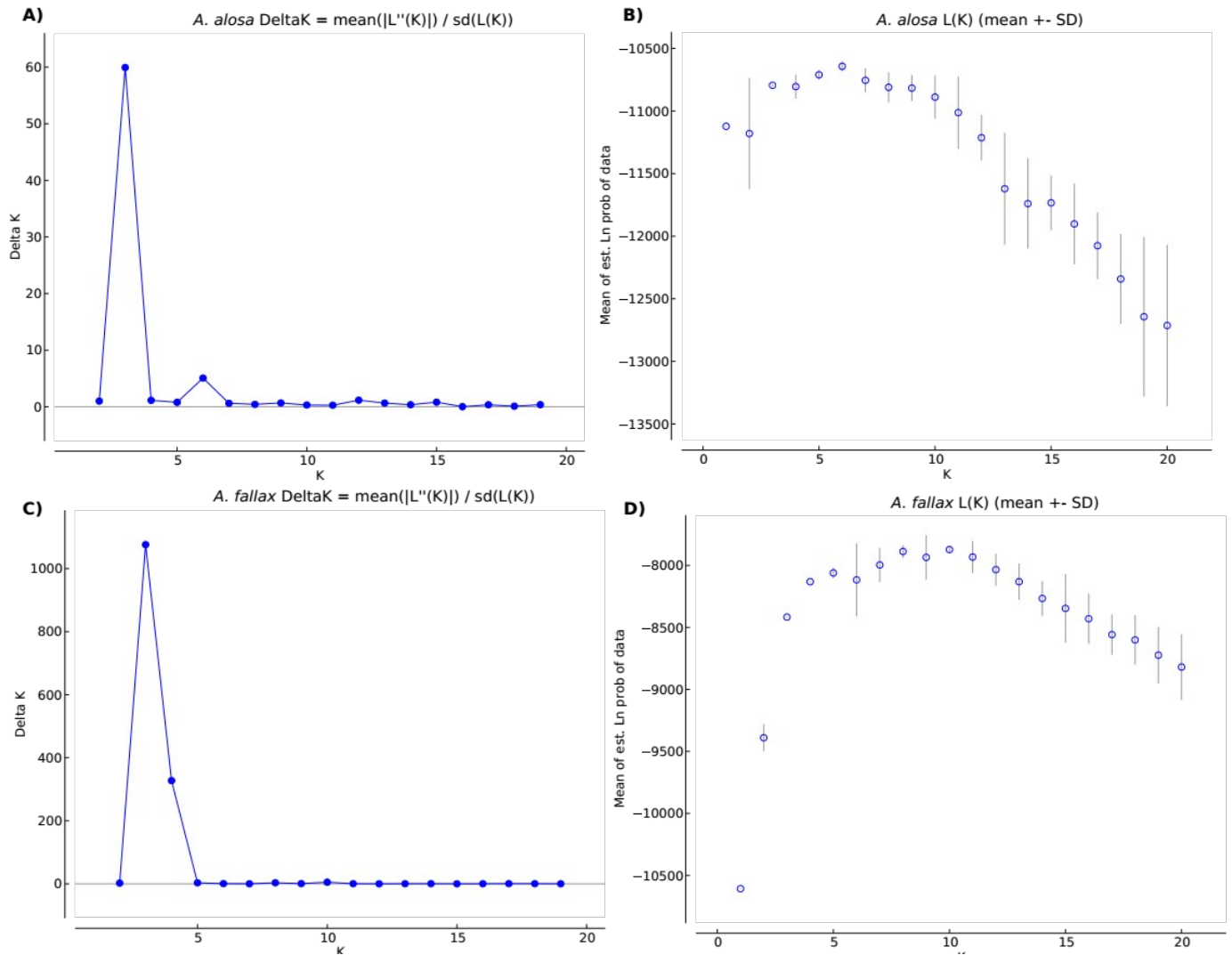

**Figure S3: A) Ln (K) and B) delta K obtained when considering *A. fallax* sampled along the Atlantic coast only and C) corresponding population genetic structure plot.**

**a) Delta K for Atlantic *A. fallax* only**

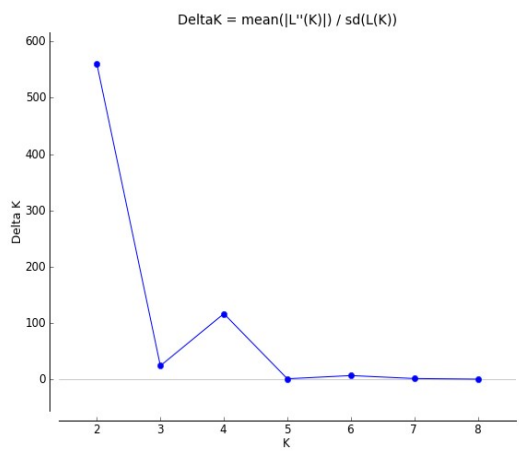

**b) Ln(K) for Atlantic *A. fallax* only**

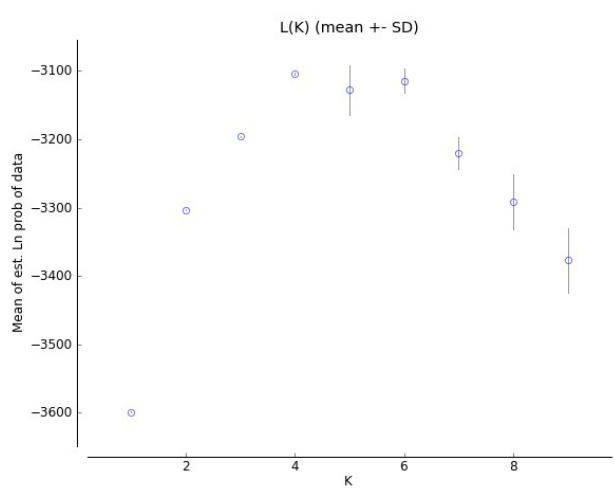

**C) Population genetic structure of *A. fallax* along the Atlantic coast. Individuals captured at sea are included.**

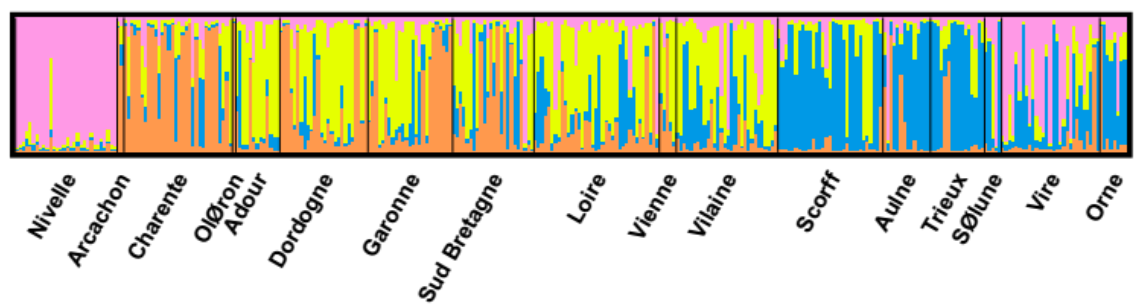

Figure S4: Bayesian Information Criteria (BIC) revealing the number of group used in the DAPC and α-score obtained in A) *A. alosa* and B) *A. fallax*.

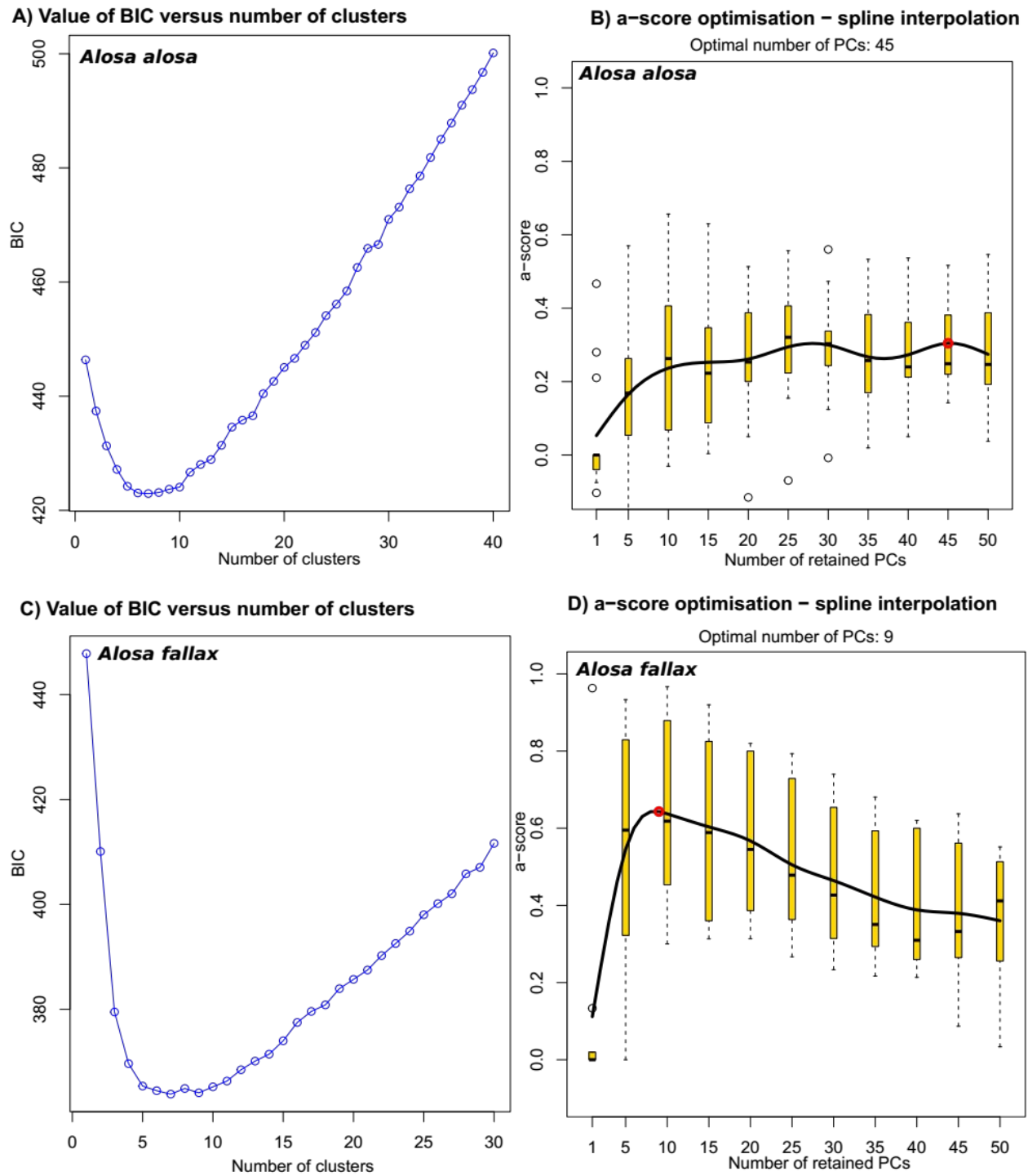

**Figure S5:** Robustness of the model choice when comparing models with ongoing gene flow (IM, SC) or ancient gene flow (AM) against a model of strict isolation (SI) in a pairwise manner

Robustness is obtained through 4,000 cross validation using the ABC model choice procedure (neural networks) on simulated data.

A) Posterior probability distribution densities of inferring the true model (M) given that the model was generated under M, i.e. $P(M | M)$. M being either SI (yellow line) or IM (black line)

B) Posterior probability of observing the IM model given it was generated under either IM or SI.

Red line = observed posterior probability on the empirical data

C) Posterior probability distribution densities of inferring the true model (M) given that the model was generated under M, i.e. $P(M | M)$. M being either SI (yellow line) or AM (black line)

D) Posterior probability of observing the AM model given it was generated under either AM or SI.

E) Posterior probability distribution densities of inferring the true model (M) given that the model was generated under M, i.e. $P(M | M)$. M being either SI (yellow line) or SC (black line)

F) Posterior probability of observing the SC model given it was generated under either SC or SI.

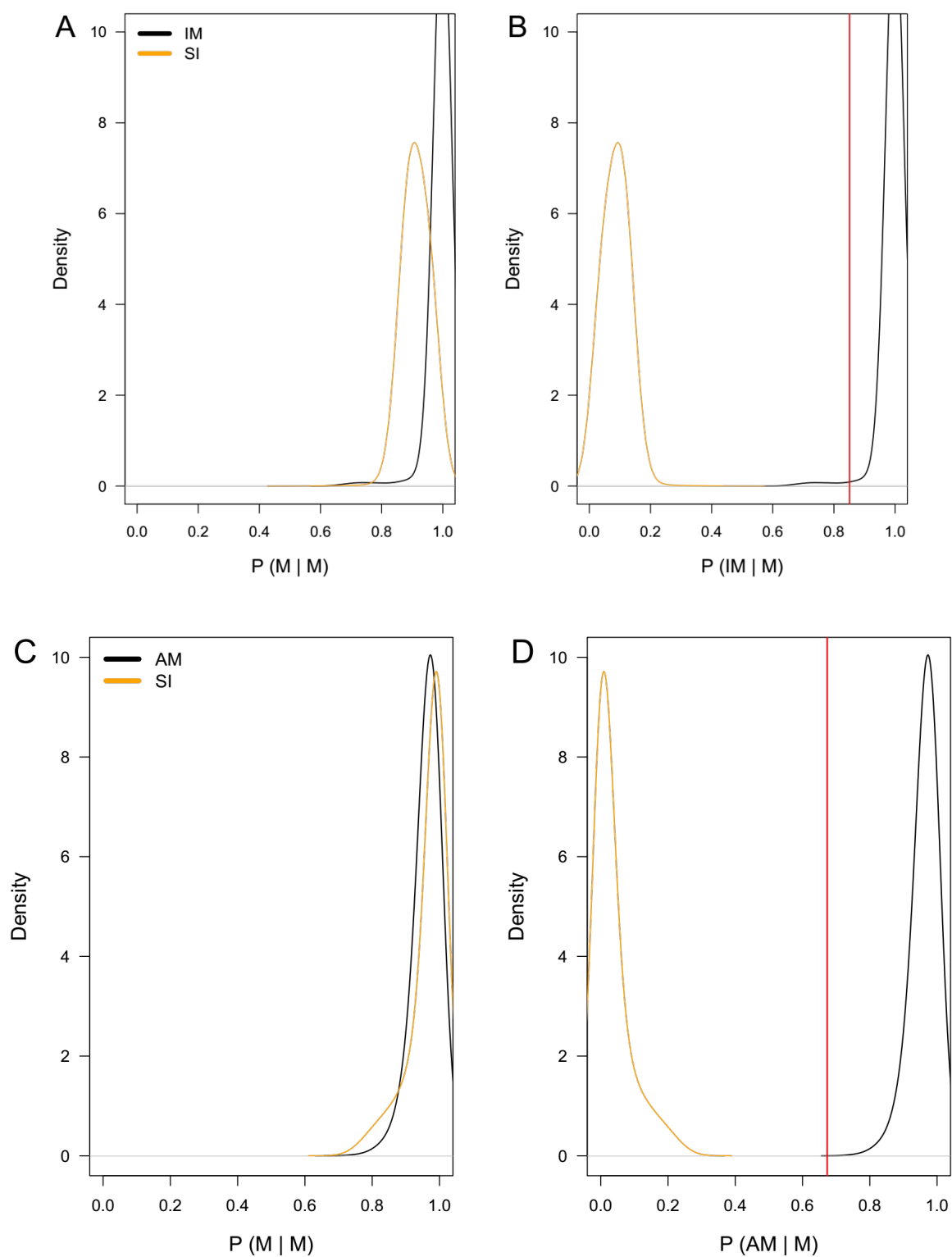

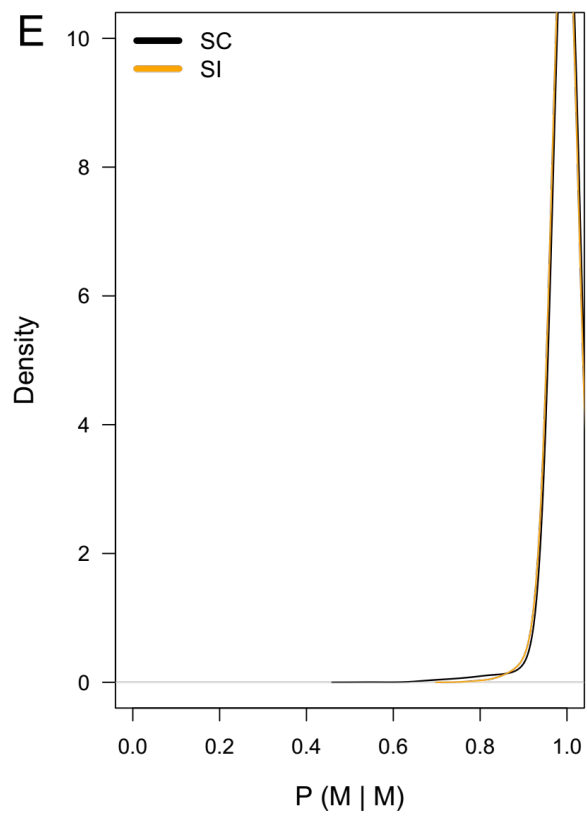

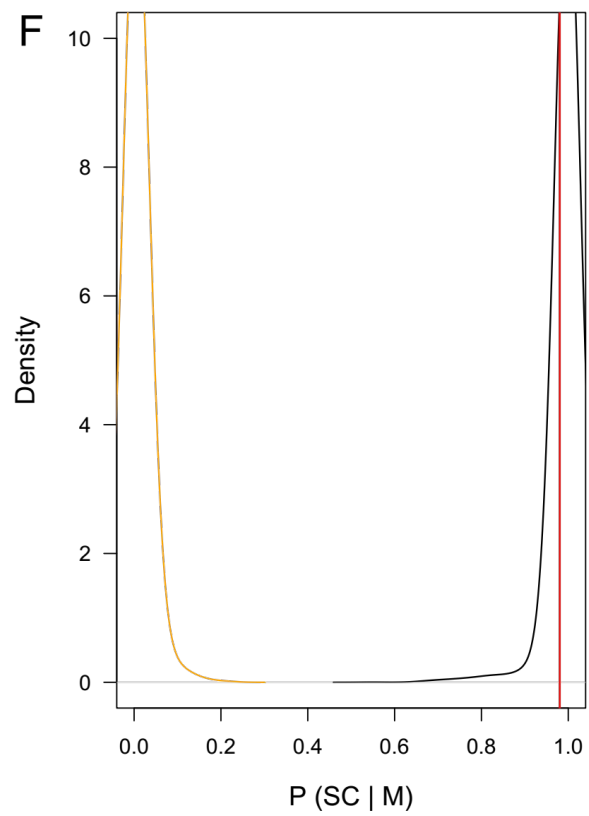

**Supplementary Figure S6: Robustness of the model choice comparing model with ongoing gene-flow (IM) against models with Ancient Gene flow (AM) or secondary Contact (SC).**

Robustness is obtained through 4,000 cross-validations using the ABC model choice procedure (neural networks) on simulated data.

A) Posterior probability distribution densities of inferring the true model (M) given that the model was generated under M, i.e. $P(M | M)$. M being either AM (yellow line) or IM (black line)

B) Posterior probability of observing the IM model given it was generated under either IM or AM.

C) Posterior probability distribution densities of inferring the true model (M) given that the model was generated under M, i.e. $P(M | M)$. M being either IM (yellow line) or SC (black line)

D) Posterior probability of observing the SC model given it was generated under either IM or SC.

Red line = posterior probability of the model observed in the empirical data.

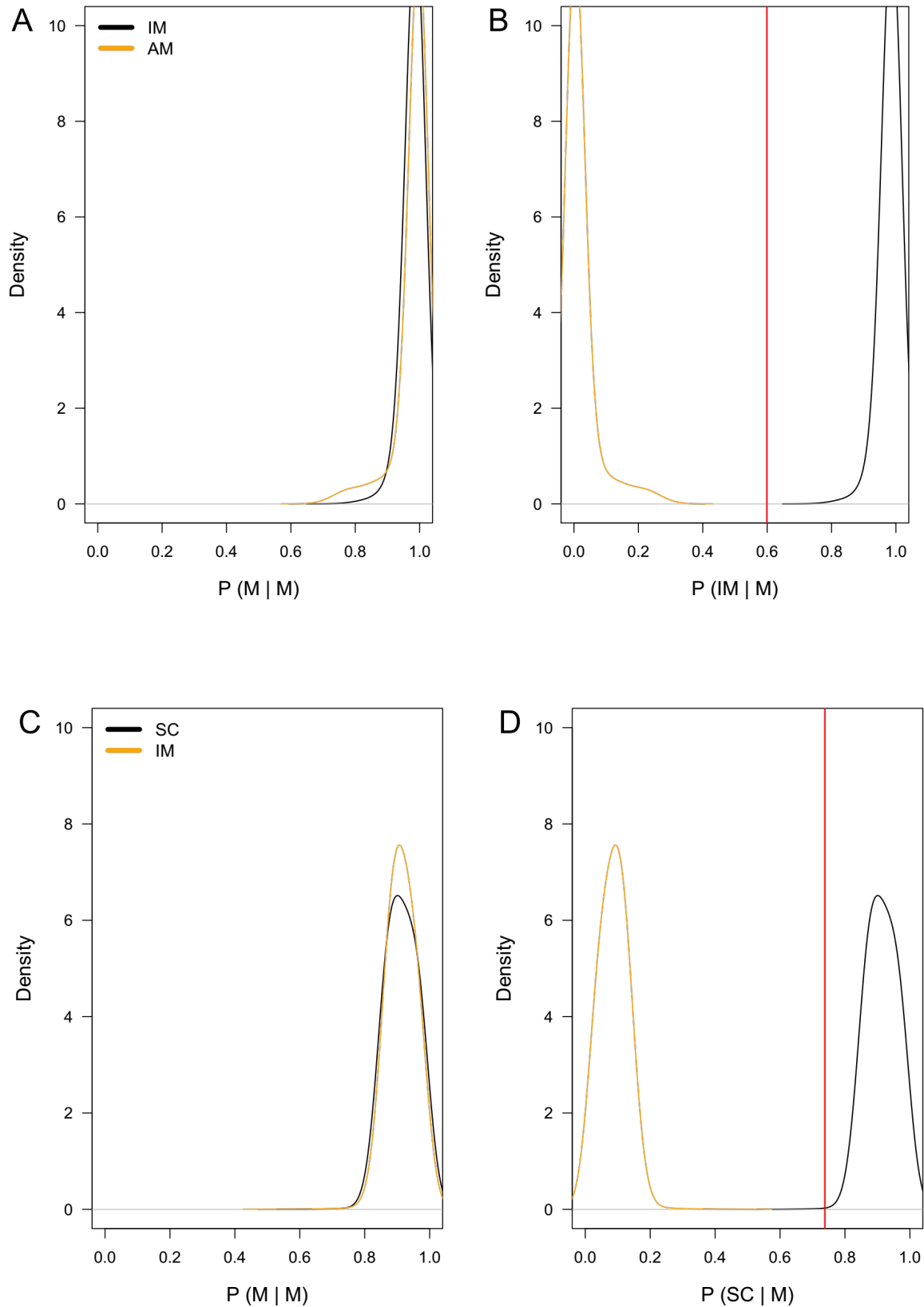

**Figure S7: Prior (grey) and Posterior (orange) distribution of parameter estimates associated to the best model inferred by ABC when considering *A. fallax* lineages:**

$\theta$ *A. fallax* Atlantic = Scaled Effective population size of *A. fallax* “Atlantic lineage”. $\theta$ *A. fallax* “Mediterranean” = Scaled Effective population size of *A. fallax* “Mediterranean lineage”. $\theta_{\text{ancestral}}$ = Effective population size of the ancestral population. All populations size are scaled by $4N_{\text{ref}}\mu$. $T_{\text{split}}$ = split time (here $T_{\text{split}} = \tau \cdot 4N_{\text{ref}}$). $T_{\text{sc}}$ = Time of Secondary Contact. $M_{12} = 4N_{\text{ref}}m_{12}$ and $M_{21} = 4N_{\text{ref}}m_{21}$ correspond to the scaled migration rate where $m_{ij}$ represents the fraction of subpopulation $i$ which is made up of migrants from subpopulation $j$ each generation. All values are provided in coalescent units and scaled by the effective reference population size ($N_{\text{ref}} = 50,000$).

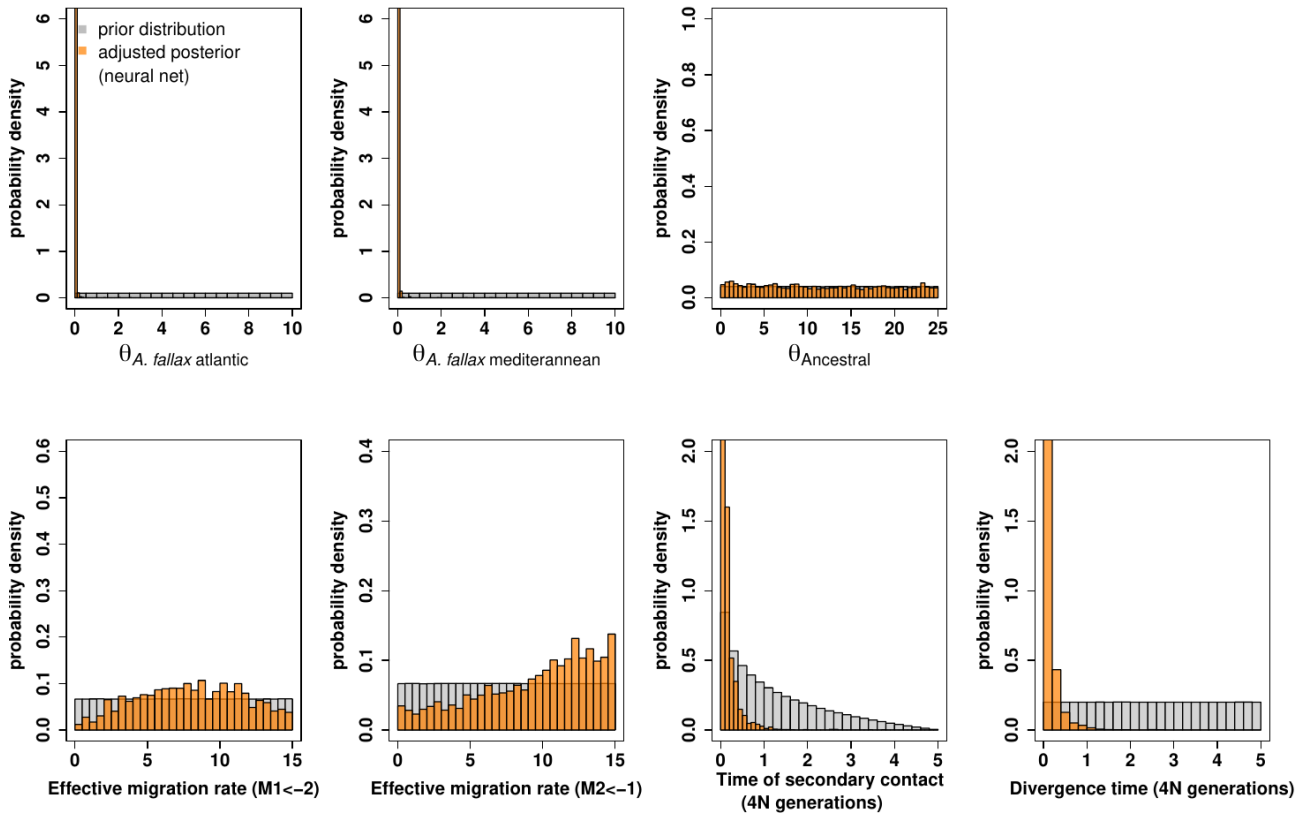

**Figure S8: Prior (grey) and Posterior (orange) distribution of parameter estimates associated to the best model inferred by ABC when considering *A. fallax* lineages within Mediterranean Sea:**

$\theta$ *A. fallax* “Mediterranean” = Scaled Effective population size of *A. fallax* “Mediterranean lineage”. $\theta$ *A. fallax* “Corsica” = Scaled Effective population size of *A. fallax* “Corsican lineage”. $\theta_{\text{ancestral}}$ = Effective population size of the ancestral population. All populations size are scaled by $4N_{\text{ref}}\mu$. Tsplit = split time (here Tsplit = $\tau \cdot 4N_{\text{ref}}$). Tsc = Time of Secondary Contact. $M12 = 4N_{\text{ref}}m12$ and $M21 = 4N_{\text{ref}}m21$ correspond to the scaled migration rate where $m_{ij}$ represents the fraction of subpopulation $i$ which is made up of migrants from subpopulation $j$ each generation. All values are provided in coalescent units and scaled by the effective reference population size ($N_{\text{ref}} = 50,000$).

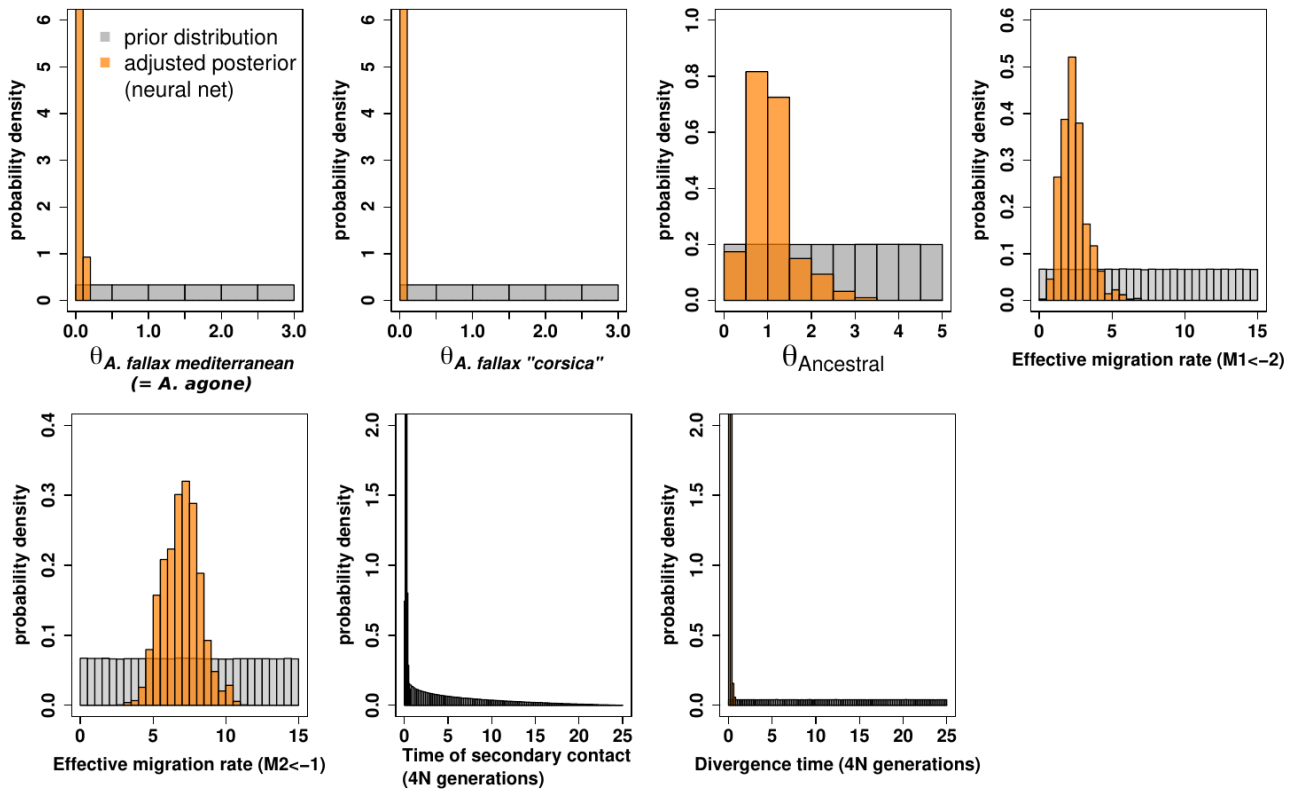
